# Image content is more important than Bouma’s Law for scene metamers

**DOI:** 10.1101/378521

**Authors:** Thomas S. A. Wallis, Christina M. Funke, Alexander S. Ecker, Leon A. Gatys, Felix A. Wichmann, Matthias Bethge

**Affiliations:** Werner Reichardt Center for Integrative Neuroscience, Eberhard Karls Universität Tübingen; Bernstein Center for Computational Neuroscience, Tübingen; Center for Neuroscience and Artificial Intelligence, Baylor College of Medicine, Houston, TX, USA; Institute for Theoretical Physics, Eberhard Karls Universität Tübingen; Presently at Apple, Inc., Cupertino, California; Neural Information Processing Group, Faculty of Science, Eberhard Karls Universität Tübingen; Max Planck Institute for Biological Cybernetics, Tübingen; First authorship shared equally.

## Abstract

We subjectively perceive our visual field with high fidelity, yet large peripheral distortions can go unnoticed and peripheral objects can be difficult to identify (crowding). A recent paper proposed a model of the mid-level ventral visual stream in which neural responses were averaged over an area of space that increased as a function of eccentricity (scaling). Human participants could not discriminate synthesised model images from each other (they were metamers) when scaling was about half the retinal eccentricity. This result implicated ventral visual area V2 and approximated “Bouma’s Law” of crowding. It has subsequently been interpreted as a link between crowding zones, receptive field scaling, and our rich perceptual experience. However, participants in this experiment never saw the original images. We find that participants can easily discriminate real and model-generated images at V2 scaling. Lower scale factors than even V1 receptive fields may be required to generate metamers. Efficiently explaining why scenes look as they do may require incorporating segmentation processes and global organisational constraints in addition to local pooling.

## Introduction

Vision science seeks to understand why things look as they do (Koffka 1935). Typically, our entire visual field looks subjectively crisp and clear. Yet our perception of the scene falling onto the peripheral retina is actually limited by at least three distinct sources: the optics of the eye, retinal sampling, and the mechanism(s) giving rise to crowding, in which our ability to identify and discriminate objects in the periphery is limited by the presence of nearby items (Bouma 1970; Pelli and Tillman 2008).^1^ Thus we can be insensitive to significant changes in the world despite our rich subjective experience.

Visual crowding has been characterised as compulsory texture perception (Parkes et al. 2001; Lettvin 1976) and compression (Balas, Nakano, and Rosenholtz 2009; Rosenholtz, Huang, and Ehinger 2012). This idea entails that we cannot perceive the precise structure of the visual world in the periphery. Rather, we are aware only of the summary statistics or ensemble properties of visual displays, such as the average size or orientation of a group of elements (Ariely 2001; Dakin and Watt 1997). One of the appeals of the summary statistic idea is that it can be directly motivated from the perspective of efficient coding: it is a form of compression. Image-computable texture summary statistics have been shown to be correlated with human performance in various tasks requiring the judgment of peripheral information, such as crowding and visual search (Rosenholtz et al. 2012; Balas, Nakano, and Rosenholtz 2009; Freeman and Simoncelli 2011; Rosenholtz 2016; Ehinger and Rosenholtz 2016). Recently, it has even been suggested that summary statistics underlie our rich phenomenal experience itself—in the absence of focussed attention, we perceive only a texture-like visual world (Cohen, Dennett, and Kanwisher 2016).

Across many tasks, summary statistic representations seem to capture aspects of peripheral vision when their pooling corresponds to “Bouma’s Law” (Rosenholtz et al. 2012; Balas, Nakano, and Rosenholtz 2009; Freeman and Simoncelli 2011; Wallis and Bex 2012; Ehinger and Rosenholtz 2016). Bouma’s Law states that objects will crowd (correspondingly, statistics will be pooled) over spatial regions corresponding to about half the retinal eccentricity (Bouma 1970; Pelli and Tillman 2008; though see Rosen, Chakravarthi, and Pelli 2014). If the visual system does indeed represent the periphery using summary statistics, then Bouma’s scaling implies that as retinal eccentricity increases, increasingly large regions of space are texturised by the visual system. If a model captured these statistics and their pooling, images could be created that are indistinguishable from the original despite being physically different (metamers). These images would be equivalent to the model and to the human visual system (Freeman and Simoncelli 2011; Wallis, Bethge, and Wichmann 2016; Portilla and Simoncelli 2000; Koenderink et al. 2017).

Freeman and Simoncelli (2011) developed a model (hereafter, FS-model) in which texture-like summary statistics were pooled over spatial regions inspired by the receptive fields in primate visual cortex. The size of neural receptive fields in ventral visual stream areas increases as a function of retinal eccentricity, and as one moves downstream from V1 to V2 and V4 at a given eccentricity. Each visual area therefore has a signature scale factor, defined as the ratio of the receptive field diameter to retinal eccentricity (Freeman and Simoncelli 2011). Similarly, the pooling regions of the FS-model also increase with retinal eccentricity with a definable scale factor. New images could be synthesised that matched the summary statistics of original images at this scale factor. As scale factor increases, texture statistics are pooled over increasingly large regions of space, resulting in more distorted synthesised images relative to the original (that is, more information is discarded).

The maximum scale factor for which the images remain indistinguishable (the critical scale) characterises perceptually-relevant compression in the visual system’s representation. If the scale factor of the model corresponded to the scaling of the visual system in the responsible visual area, and information in upstream areas was irretrievably lost, then the images synthesised by the model should be indistinguishable while discarding as much information as possible. That is, we seek the maximum compression that is perceptually lossless. Larger scale factors would discard more information than the relevant visual area and therefore the images should look different. Smaller scale factors preserve information that could be discarded without any perceptual effect.

Crucially, it is the *minimum* critical scale over images that is important for the scaling theory. If the visual system computes summary statistics over fixed (image-independent) pooling regions in the same way as the model, then the model must be able to produce metamers for all images. While images may vary in their individual critical scales, the image with the smallest critical scale determines the maximum compression for appearance to be matched in general.

Freeman and Simoncelli showed that the largest scale factor for which *two synthesised images* could not be told apart was 0.5, or pooling regions of about half the eccentricity. This scaling matched the signature of area V2, and also matched the approximate value of Bouma’s Law. Subsequently, this result has been interpreted as a link between receptive field scaling, crowding, and our rich phenomenal experience (e.g. Block 2013, Cohen, Dennett, and Kanwisher (2016), Landy (2013), Movshon and Simoncelli (2014), Seth (2014)). These interpretations imply that the FS-model creates metamers *for natural scenes*. However, observers in Freeman and Simoncelli’s experiment never saw the original scenes, but only compared synthesised images to each other. Showing that two model samples are indiscriminable from each other could yield trivial results. For example, two white noise samples matched to the mean and contrast of a natural scene would be easy to discriminate from the scene but hard to discriminate from each other. Wallis, Bethge and Wichmann (2016) showed that observers could easily discriminate Portilla and Simoncelli (2000) textures from original images in the periphery, but did not test the FS-model. The Portilla and Simoncelli model makes no explicit connection to neural receptive field scaling. In addition, relative to the textures tested by Wallis et al., the pooling region overlap used in the FS-model provides a strong constraint on the resulting syntheses, making the images much more similar to the originals. It is therefore entirely possible that the FS-model produces metamers for natural scenes for scale factors of 0.5. Here we test this, and compare the results to our own model using CNN texture features.

## Results

We tested whether the FS-model can produce metamers using an oddity design in which the observer had to pick the odd image out of three successively shown images (Fig 1E). In a 3-alternative oddity paradigm, performance for metamerism would lie at 0.33 (dashed horizontal line, Figure 1F). We used two comparison conditions: either observers compared two model syntheses to each other (synth vs synth; as in Freeman and Simoncelli 2011) or the original image to a model synthesis (orig vs synth). As in the original paper (Freeman and Simoncelli 2011) we measured the performance of human observers for images synthesised with different scale factors (using Freeman and Simoncelli’s code, see Methods). To quantify the critical scale factor we fit the same nonlinear model as Freeman and Simoncelli, which parameterises sensitivity as a function of critical scale and gain, as a mixed-effects model with random effects of participant and image (see Methods).

In previous experiments (see Supplementary Figures S8 and S9), we observed that texture-like distortions are more visible when they fall over image regions containing inhomogenous structure, long edges, or borders between different surfaces than when they fall into texture-like regions. We therefore explicitly compared “scene-like” images containing inhomogenous structures to “texture-like” images containing more homogenous or periodically-patterned content in the periphery. We hand-selected ten images from each class (Figure 1A and Figure 1B)^2^.

When participants compared synthesised images to each other as in Freeman and Simoncelli (Figure 1F, synth vs synth), there was little evidence that the critical scale depended on the image content (scene-vs texture-like). The difference in critical scale between texture-like and scene-like images was 0.09, 95% CI [-0.03, 0.24], p(*β* < 0) = 0.078. This result is consistent with Freeman and Simoncelli, who reported no dependency on image. It seems likely that this is because comparing synthesised images to each other means that the model has removed higher-order structure that might allow discrimination. All images appear distorted, and the task becomes one of identifying a specific distortion pattern. While the critical scales we find are somewhat lower than those reported by Freeman and Simoncelli (2011; Figure 1G), they are within the range of other reported critical scale factors (Freeman and Simoncelli 2013). One striking difference between our results for synth vs synth and those of Freeman and Simoncelli is that the performance of our participants was quite poor even for large scale factors. This may be because because we used more images in our experiment than Freeman and Simoncelli, so participants were less familiar with the distortions that could appear. We have replicated these results in an experiment using the same ABX task as in Freeman and Simoncelli (Figure S5).

Comparing the original image to model syntheses yielded a different pattern of results. First, participants are able to discriminate the original images from their FS-model syntheses at scale factors of 0.5 (Figure 1F). Performance lies well above chance for all participants: these images are not metamers. This holds for both scene-like and texture-like images. Furthermore, critical scale depends on the image type. Model syntheses match the texture-like images on average with scale factors of approximately 0.25 (the smallest value we could generate using the FS-model). In contrast, the scene-like images are quite discriminable from their model syntheses at this scale. Correspondingly, texture-like images had higher critical scales than scene-like images on average (0.13, 95% CI [0.06, 0.22], p(*β* < 0) = 0.001). Thus, smaller pooling regions are required to make metamers for scene-like images than for texture-like images.

As noted above, the image with the minimum critical scale determines the largest compression that can be applied for the scaling model to hold. For two images (Figure 2A and E) the nonlinear mixed-effects model estimated critical scales of approximately 0.14 (see Figure 1G, diamonds). However, examining the individual data for these images (Figure 2D and H) reveals that these critical scale estimates are largely determined by the hierarchical nature of the mixed-effects model, not the data itself. Both images remain highly discriminable for the lowest scale factor we could generate. This suggests that the mixed-effects model critical scale may be an overestimate of the true scale factor required to generate metamers. Thus, human observers are highly sensitive to the relatively small distortions produced by the FS-model at scale factors of 0.25 (compare Figure 2B and F at scale 0.25 and C and G at scale 0.46 to images A and B).

## Discussion

It is a popular idea that the appearance of scenes in the periphery is described by summary statistic textures captured at the scaling of V2 neural populations. In contrast, here we show that humans are very sensitive to the difference between original and model-matched images at this scale (Figure 1). A recent preprint (Deza, Jonnalagadda, and Eckstein 2017) finds a similar result in a set of 50 images, and our results are also consistent with the speculations made by Wallis et al. based on their experiments with Portilla and Simoncelli textures (Wallis, Bethge, and Wichmann 2016). Together, these results show that the pooling of texture-like features in the FS-model at the scaling of V2 receptive fields does not explain the appearance of natural images.

If the peripheral appearance of visual scenes is explained by image-independent pooling of texture-like features, then the pooling regions must be small. Consider that participants in our experiment could easily discriminate the images in Figure 2B and F from those in Figure 2A and E respectively. Therefore, images synthesised at a truly metameric scaling must remain extremely close to the original: at least as small as V1 neurons, and likely even lower (Figure 2). This may even be consistent with scaling in precortical visual areas. For example, the scaling of retinal ganglion cell receptive fields at the average eccentricity of our stimuli (6 degrees) is approximately 0.08 for the surround (Croner and Kaplan 1995) and 0.009 for the centre (Dacey and Petersen 1992). It becomes questionable how much is learned about compression in the visual system using such an approach, beyond the aforementioned, relatively well-studied limits of optics and retinal sampling (e.g. Wandell 1995; Watson 2014).

Furthermore, it can be seen via simple demonstration that we can be quite insensitive to even large texture-like distortions, so long as these fall on texture-like regions of the input image. The “China Lane” sign in Figure 3A has been distorted in B (using texture features from Gatys et al (2015)). The same type of distortion in a texture-like region of the image is far less visible (the brickwork in the image centre; FS-model result Figure 3C; see also Figure S13). It is the image content, not retinal eccentricity, that is the primary determinant of the visibility of at least some summary statistic distortions. Requiring information to be preserved at V1 or smaller scaling would therefore be rather inefficient from the standpoint of compression: small scale factors will preserve texture-like structure that could be compressed without affecting appearance.

It may seem trivial that a texture statistic model better captures the appearance of textures than non-textures. However, if the human visual system represents the periphery as a set of textures, and these models are sufficient approximations of this representation, then image content should not matter— because scene-like retinal inputs in the periphery are transformed into textures by the visual system.

Perhaps the texture scaling theory might hold at larger scales but the FS-model texture features themselves are insufficient to capture natural scene appearance. To test whether improved texture features (Gatys, Ecker, and Bethge 2015; Wallis et al. 2017) could help in matching appearance for scenes, we developed a new model (CNN-model) that was inspired by the FS-model but uses the texture features of a convolutional neural network (see Methods and Figures S6–9). The CNN-model shows very similar behaviour to the FS-model such that human performance for scene-like images is higher than for texture-like images (triangles in Figure 1D and Figure 2), and the CNN-model also fails to create metamers for all images (see also Figures S9, S12–13). Furthermore, the NeuroFovea model of Deza et al (2017), which like our CNN-model uses deep neural network texture features, also fails to capture scene appearance. Together, these results show that no known summary statistic pooling model is sufficient to match the appearance of arbitrary natural scenes at computationally feasible scale factors.

What, then, is the missing ingredient that could capture appearance while compressing as much information as possible? Through the Gestalt tradition, it has long been known that the appearance of local image elements can crucially depend on the context in which they are placed. For example, Saarela et al (2009) and Manassi et al (2013) found that global stimulus configuration modulates crowding (see also Vickery et al. 2009), and the results of Neri (2017) suggest that early global segmentation processes influence local perceptual sensitivity. Potentially, global scene organisation needs to be considered if one wants to capture appearance—yet current models that texturise local regions do not explicitly include perceptual organisation (Herzog et al. 2015). We speculate that segmentation and grouping processes are critical for efficiently matching scene appearance, and therefore the approach of uniformly computing summary statistics without including these processes will require preserving much of the original image structure by making pooling regions very small. A parsimonious model capable of compressing as much information as possible might need to adapt either the size and arrangement of pooling regions or the feature representations to the image content.

**Figure 1.**
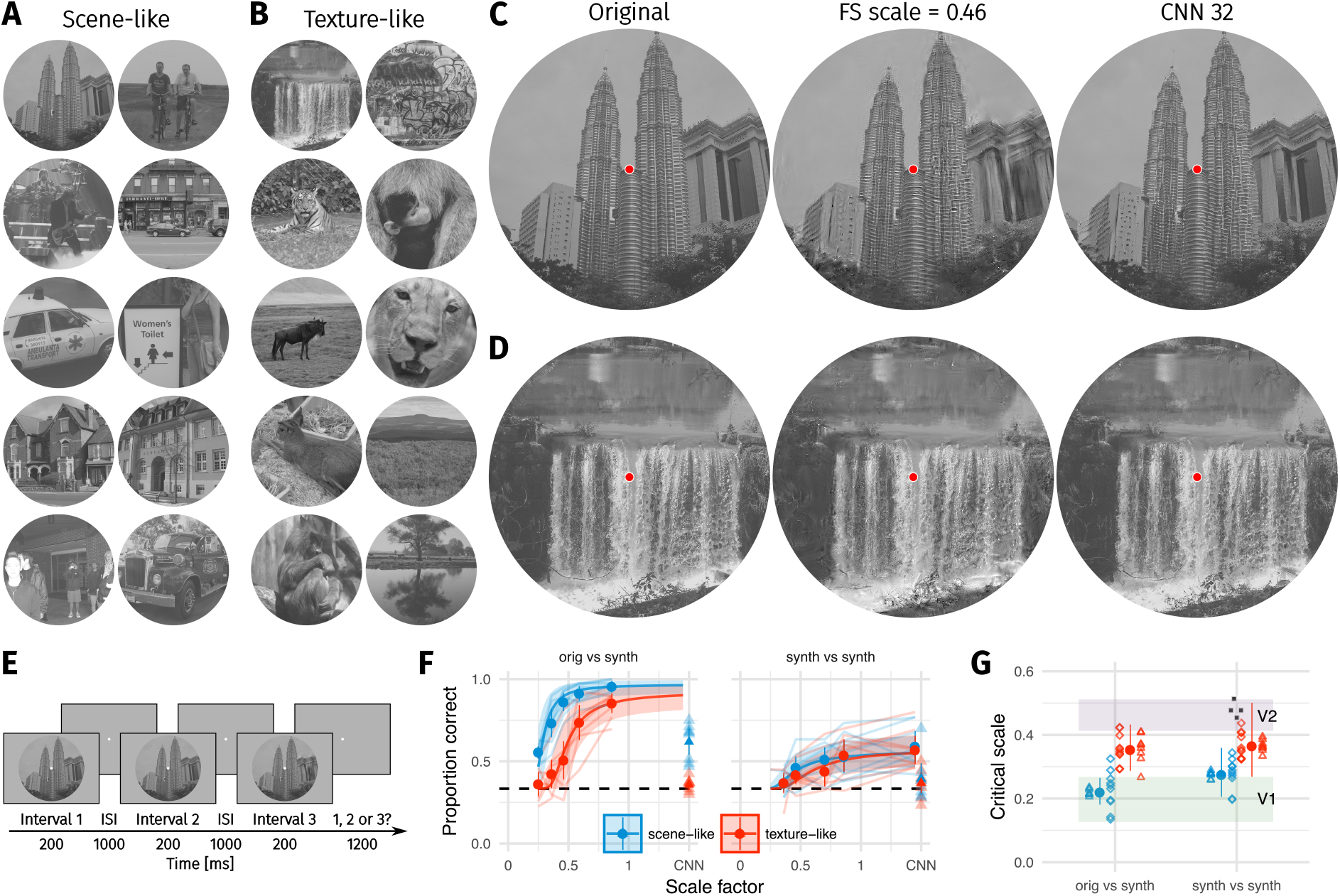
Two texture pooling models fail to match arbitrary scene appearance. We selected ten scene-like (**A**) and ten texture-like (**B**) images and synthesised images to match them using the Freeman & Simoncelli model (FS scale 0.46 shown) or a model using CNN texture features (CNN 32; example scene and texture-like stimuli shown in C and D respectively). **E**: The oddity paradigm. Three images were presented in sequence, with two being physically-identical and one being the oddball. Participants indicated which image was the oddball (1, 2 or 3). On ”orig vs synth” trials participants compared real and synthesised images, whereas on ”synth vs synth” trials participants compared two images synthesised from the same model. **F**: Performance as a function of scale factor (pooling region diameter divided by eccentricity) in the Freeman-Simoncelli model (circles) and for the CNN 32 model (triangles; arbitrary x-axis location). Points show grand mean ±2 SE over participants; faint lines link individual participant performance levels (FS-model) and faint triangles show individual CNN 32 performance. Solid curves and shaded regions show the fit of a nonlinear mixed-effects model estimating the critical scale and gain. Participants are still above chance for scene-like images in the original vs synth condition for the lowest scale factor of the FS-model we could generate, and for the CNN 32 model, indicating that neither model succeeds in producing metamers. **G**: When comparing original and synthesised images, estimated critical scales (scale at which performance rises above chance) are lower for scene-like than for texture-like images. Points with error bars show population mean and 95% credible intervals. Triangles show posterior means for participants; diamonds show posterior means for images. Black squares show critical scale estimates of the four participants from Freeman & Simoncelli reproduced from that paper (x-position jittered to reduce overplotting); shaded regions denote the receptive field scaling of V1 and V2 estimated by Freeman & Simoncelli.

**Figure 2.**
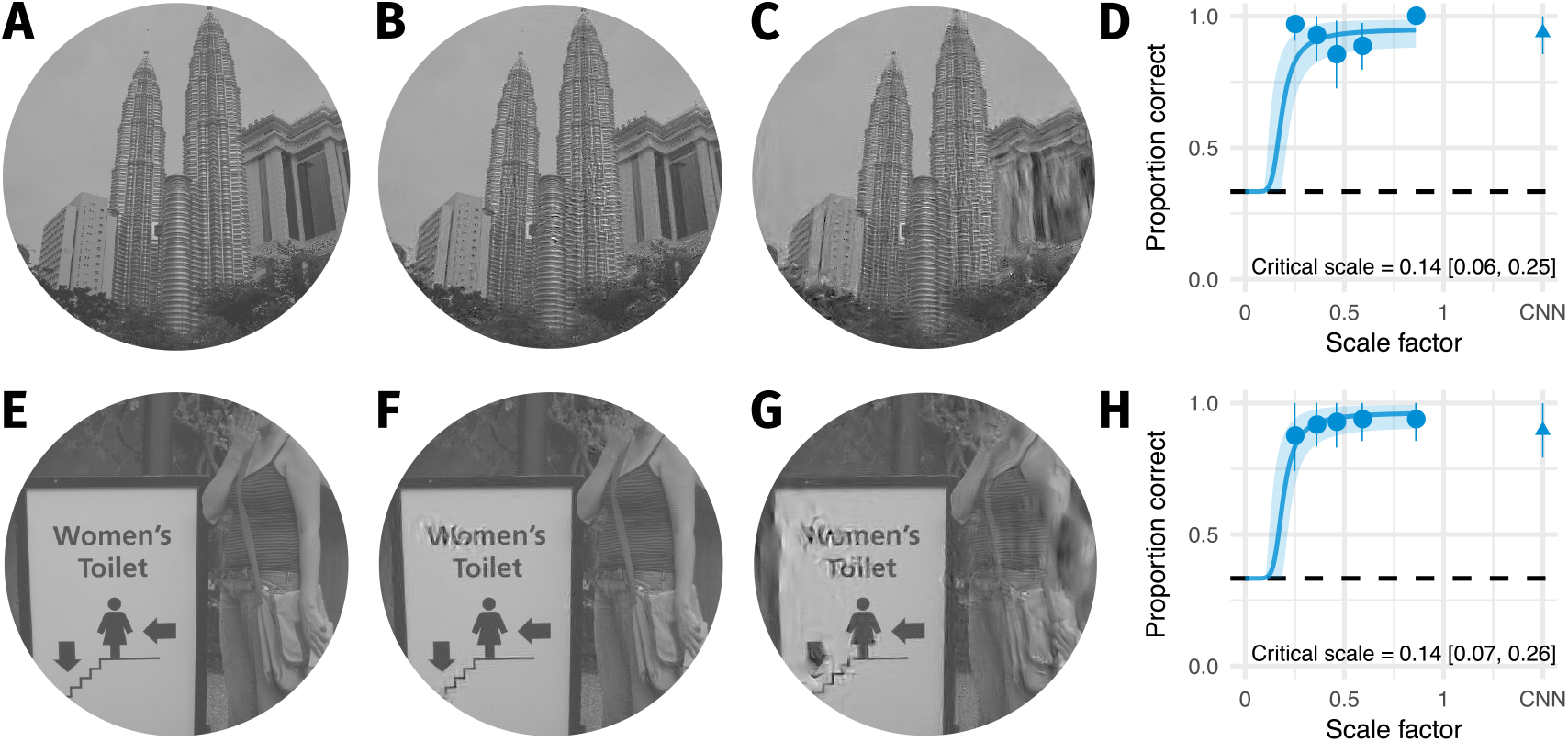
The two images with smallest critical scale estimates are highly discriminable even for the lowest scale factor we could generate. **A**: The original image. **B**: An example FS synthesis at scale factor 0.25. **C**: An example FS synthesis at scale factor 0.46. **D**: The average data for this image. Points and error bars show grand mean and ±2 SE over participants, solid curve and shaded area show posterior mean and 95% credible intervals from the mixed-effects model. Embedded text shows posterior mean and 95% credible interval on the critical scale estimate for this image. **E–G**: Same as A–D for the image with the second-lowest critical scale. Note that in both cases the model is likely to overestimate critical scale.

Our results do not undermine the considerable empirical support for the periphery-as-summary-statistic theory as a description of visual performance. Humans can judge summary statistics of visual displays (Ariely 2001; Dakin and Watt 1997), summary statistics can influence judgments where other information is lost (Fischer and Whitney 2011; Faivre, Berthet, and Kouider 2012), and the information preserved by summary statistic stimuli may offer an explanation for performance in various visual tasks (Rosenholtz et al. 2012; Balas, Nakano, and Rosenholtz 2009; Rosenholtz, Huang, and Ehinger 2012; Keshvari and Rosenholtz 2016; Chang and Rosenholtz 2016; Zhang et al. 2015; Whitney, Haberman, and Sweeny 2014; Long et al. 2016; though see Agaoglu and Chung 2016; Herzog et al. 2015; Francis, Manassi, and Herzog 2017). Texture-like statistics may even provide the primitives—appropriately organised on a global scale—from which form is constructed (Lettvin 1976). However, one additional point merits further discussion. The studies by Rosenholtz and colleagues primarily test summary statistic representations by showing that performance with summary statistic stimuli viewed foveally is correlated with peripheral performance with real stimuli. This means that the summary statistic preserves sufficient information to explain the performance of tasks in the periphery. Our results show that these summary statistics are insufficient to match scene appearance, at least under the pooling scheme used in the Freeman and Simoncelli model at computationally feasible scales. This shows the usefulness of scene appearance matching as a test: a parsimonious model that matches scene appearance would be expected to also preserve enough information to show correlations with peripheral task performance; the converse does not hold.

While it may be useful to consider summary statistic pooling in accounts of visual performance, to say that summary statistics can account for phenomenological experience of the visual periphery (Cohen, Dennett, and Kanwisher 2016; see also Block 2013; Seth 2014) seems premature in light of our results (see also Haun et al. 2017). Cohen et al (2016) additionally posit that focussed spatial attention can in some cases overcome the limitations imposed by a summary statistic representation. We instead find little evidence that participants’ ability to discriminate real from synthesised images is improved by cueing spatial attention, at least in our experimental paradigm and for our CNN-model (Figure S11).

**Figure 3.**
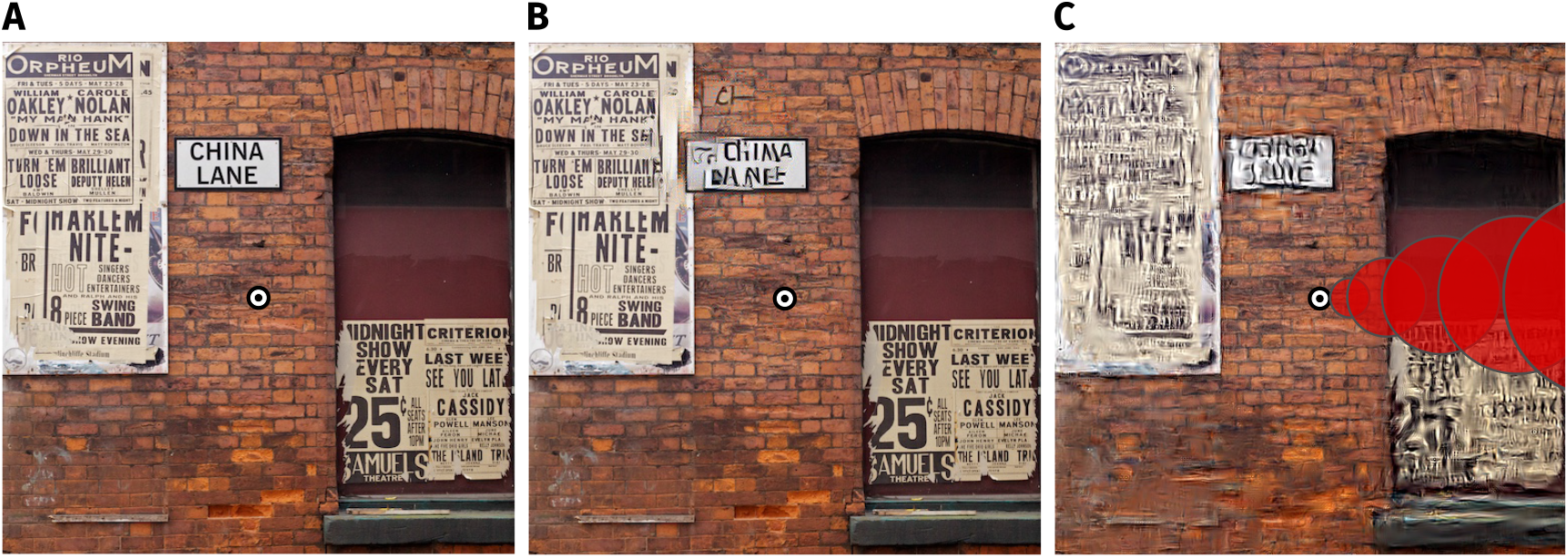
The visibility of texture-like distortions depends on image content. **A**: ”Geotemporal Anomaly” by Pete Birkinshaw (2010; re-used under a CC-BY 2.0 license). **B**: Two texture-like distortions have been introduced into circular regions of the scene in A (see Figure S1 for higher resolution). The distortion in the upper-left is quite visible, even with central fixation on the bullseye, because it breaks up the high-contrast contours of the text. The second distortion occurs on the brickwork centered on the bullseye, and is more difficult to see (you may not have noticed it until reading this caption). The visibility of texture-like distortions can depend more on image content than on retinal eccentricity (see also Figure S13). **C**: Results synthesised from the FS-model at scale 0.46 for comparison. Pooling regions depicted for one angular meridian as overlapping red circles; real pooling regions are smooth functions tiling the whole image. Pooling in this fashion reduces large distortions compared to B, but our results show that this is insufficient to match appearance.

One exciting aspect of Freeman and Simoncelli (2011) was the promise of inferring a critical brain region via a receptive field size prediction derived from psychophysics. Indeed, aspects of this promise have since received empirical support: the presence of texture-like features can discriminate V2 neurons from V1 neurons (Freeman et al. 2013; Ziemba et al. 2016; see also Okazawa, Tajima, and Komatsu 2015). Discarding all higher-order structure not captured by the candidate model by comparing syntheses to each other, thereby isolating only features that change, may therefore be a useful way to distinguish sequential feedforward processing stages in neurons. On the other hand, explaining appearance is—to return to Koffka—a grand goal of vision science. For this the original vs synthesised comparison is key. Our results suggest that it is wrong to believe that “appearance”, even only peripheral appearance, can be tied solely to the scaling of receptive fields in any single brain region or to Bouma’s Law.

## Methods

All stimuli, data and code to reproduce the figures and statistics reported in this paper are available at http://dx.doi.org/10.5281/zenodo.1475112. This document was prepared using the knitr package (Xie 2013, 2015) in the R statistical environment (R Core Team 2017; Wickham and Francois 2016; Wickham 2009, 2011; Auguie 2016; Arnold 2016) to improve its reproducibility.

### Participants

Eight observers participated in the experiment: authors CF and TW, a research assistant unfamiliar with the experimental hypotheses, and five näıve participants recruited from an online advertisement pool who were paid 10 Euro / hr for two one-hour sessions. An additional näıve participant was recruited but showed insufficient eyetracking accuracy (see below). All participants signed a consent form prior to participating. Participants reported normal or corrected-to-normal visual acuity. All procedures conformed to Standard 8 of the American Psychological Association’s “Ethical Principles of Psychologists and Code of Conduct” (2010).

### Stimuli

We selected 10 “scene-like” and 10 “texture-like” source images from the MIT 1003 scene dataset (Judd, Durand, and Torralba 2012; Judd et al. 2009). We used images from this dataset to allow better comparison to related experiments (see Supplementary Material). A square was cropped from the center of the original image and downsampled to 512 x 512 px. The images were converted to grayscale and standardized to have a mean gray value of 0.5 (scaled [0,1]) and an RMS contrast (σ/µ) of 0.3.

#### Freeman and Simoncelli syntheses

We synthesised images using the FS-model (Freeman and Simoncelli 2011, code available from https://github.com/freeman-lab/metamers). Four unique syntheses were created for each source image at each of eight scale factors (0.25, 0.36, 0.46, 0.59, 0.7, 0.86, 1.09, 1.45), using 50 gradient steps as in Freeman and Simoncelli. Pilot experiments with stimuli generated with 100 gradient steps produced similar results. To successfully synthesise images at scale factors of 0.25 and 0.36 it was necessary to increase the central region of the image in which the original pixels were perfectly preserved (pooling regions near the fovea become too small to compute correlation matrices). Scales of 0.25 used a central radius of 32 px (0.8 dva in our viewing conditions) and scales 0.36 used 16 px (0.4 dva). This change should, if anything, make syntheses even harder to discriminate from the original image. All other parameters of the model were as in Freeman and Simoncelli. Synthesising an image with scale factor 0.25 took approximately 35 hours, making a larger set of syntheses or source images infeasible. It was not possible to reliably generate images with scale factors lower than 0.25 using the code above.

#### CNN model syntheses

The CNN pooling model (triangles in Figure 1) was inspired by the model of Freeman and Simoncelli, with two primary differences: first, we replaced the Portilla and Simoncelli (2000) texture features with the texture features derived from a convolutional neural network (Gatys, Ecker, and Bethge 2015), and second, we simplified the “foveated” pooling scheme for computational reasons. Specifically, for the CNN 32 model presented above, the image was divided up into 32 angular regions and 28 radial regions, spanning the outer border of the image and an inner radius of 64 px. Within each of these regions we computed the mean activation of the feature maps from a subset of the VGG-19 network layers (conv1 1, conv2 1, conv3 1, conv4 1, conv5 1). To better capture long-range correlations in image structure, we computed these radial and angular regions over three spatial scales, by computing three networks over input sizes 128, 256 and 512 px. Using this multiscale radial and angular pooling representation of an image, we synthesised new images to match the representation of the original image via iterative gradient descent (Gatys, Ecker, and Bethge 2015). Specifically, we minimised the mean-squared distance between the original and a target image, starting from Gaussian noise outside the central 64 px region, using the L-BFGS optimiser as implemented in scipy (Jones, Oliphant, and Peterson 2001) for 1000 gradient steps, which we found in pilot experiments was sufficient to produce small (but not zero) loss. Further details, including tests of other variants of this model, are provided in the Supplement.

### Equipment

Stimuli were displayed on a VIEWPixx 3D LCD (VPIXX Technologies Inc., Saint-Bruno-de-Montarville, Canada; spatial resolution 1920 x 1080 pixels, temporal resolution 120 Hz, operating with the scanning backlight turned off in normal colour mode). Outside the stimulus image the monitor was set to mean grey. Participants viewed the display from 57 cm (maintained via a chinrest) in a darkened chamber. At this distance, pixels subtended approximately 0.025 degrees on average (approximately 40 pixels per degree of visual angle). The monitor was linearised (maximum luminance 260 cd/m^2^) using a Konica-Minolta LS-100 (Konica-Minolta Inc., Tokyo, Japan). Stimulus presentation and data collection was controlled via a desktop computer (Intel Core i5-4460 CPU, AMD Radeon R9 380 GPU) running Ubuntu Linux (16.04 LTS), using the Psychtoolbox Library (version 3.0.12, Brainard 1997; Kleiner, Brainard, and Pelli 2007; Pelli 1997), the Eyelink toolbox (Cornelissen, Peters, and Palmer 2002) and our internal iShow library (http://dx.doi.org/10.5281/zenodo.34217) under MATLAB (The Mathworks Inc., Natick MA, USA; R2015b). Participants’ gaze position was monitored by an Eyelink 1000 (SR Research) video-based eyetracker.

### Procedure

On each trial participants were shown three images in succession; two images were identical, one image was different (the “oddball”, which could occur first, second or third with equal probability). The oddball could be either a synthesised or a natural image (in the orig vs synth condition; counterbalanced), whereas the other two images were physically the same as each other and from the opposite class as the oddball. In the synth vs synth condition (as used in Freeman and Simoncelli), both oddball and foil images were (physically different) model synths. The participant identified the temporal position of the oddball image via button press. Participants were told to fixate on a central point (Thaler et al. 2013) presented in the center of the screen. The images were centred around this spot and displayed with a radius of 512 pixels (i.e. images were upsampled by a factor of two for display), subtending ≈ 12.8° at the eye. Images were windowed by a circular cosine, ramping the contrast to zero in the space of 52 pixels. The stimuli were presented for 200 ms, with an inter-stimulus interval of 1000 ms (making it unlikely participants could use motion cues to detect changes), followed by a 1200 ms response window. Feedback was provided by a 100 ms change in fixation cross brightness. Gaze position was recorded during the trial. If the participant moved the eye more than 1.5 degrees away from the fixation spot, the trial immediately ended and no response was recorded; participants saw a feedback signal (sad face image) indicating a fixation break. Prior to the next trial, the state of the participant’s eye position was monitored for 50 ms; if the eye position was reported as more than 1.5 degrees away from the fixation spot a recalibration was triggered. The inter-trial interval was 400 ms.

Scene-like and texture-like images were compared under two comparison conditions (orig vs synth and synth vs synth; see main text). Image types and scale factors were randomly interleaved within a block of trials (with a minimum of one trial from another image in between) whereas comparison condition was blocked. Participants first practiced the task and fixation control in the orig vs synth comparison condition (scales 0.7, 0.86 and 1.45); the same images used in the experiment were also used in practice to familiarise participants with the images. Participants performed at least 60 practice trials, and were required to achieve at least 50% correct responses and fewer than 20% fixation breaks before proceeding (as noted above, one participant failed). Following successful practice, participants performed one block of orig vs synth trials, which consisted of five FS-model scale factors (0.25, 0.36, 0.46, 0.59, 0.86) plus the CNN 32 model, repeated once for each image to give a total of 120 trials. The participant then practiced the synth vs synth condition for at least one block (30 trials), before continuing to a normal synth vs synth block (120 trials; scale factors of 0.36, 0.46, 0.7, 0.86, 1.45). Over two one-hour sessions, näıve participants completed a total of four blocks of each comparison condition in alternating order (except for one participant who ran out of time to complete the final block). Authors performed more blocks (total 11).

### Data analysis

We discarded trials for which participants made no response (N = 66) and broke fixation (N = 239), leaving a total of 7555 trials for further analysis. To quantify the critical scale as a function of the scale factor s, we used the same 2-parameter function for discriminability *d*′ fitted by Freeman and Simoncelli:

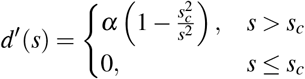

consisting of the critical scale *s_c_* (below which the participant cannot discriminate the stimuli) and a gain parameter a (asymptotic performance level in units of *d*′). This *d*′ value was transformed to proportion correct using a Weibull function as in Wallis et al (2016):

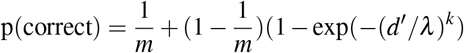

with *m* set to three (the number of alternatives), and scale *λ* and shape *k* parameters chosen by minimising the squared difference between the Weibull and simulated results for oddity as in Craven (1992). The posterior distribution over model parameters (*s_c_* and *α*) was estimated in a nonlinear mixed-effects model with fixed effects for the experimental conditions (comparison and image type) and random effects for participant (crossed with comparison and image type) and image (crossed with comparison, nested within image type), assuming binomial variability. Estimates were obtained by a Markov Chain Monte Carlo (MCMC) procedure implemented in the Stan language (version 2.16.2, Stan Development Team 2017; Hoffman and Gelman 2014), with the model wrapper package brms (version 1.10.2, Bürkner 2017) in the R statistical environment. The model parameters were given weakly-informative prior distributions, which provide information about the plausible scale of parameters but do not bias the direction of inference. Specifically, both critical scale and gain were estimated on the natural logarithmic scale; the mean log critical scale (intercept) was given a Gaussian distribution prior with mean -0.69 (corresponding to a critical scale of approximately 0.5—i.e. centred on the result from Freeman and Simoncelli) and sd 1, other fixed-effect coefficients were given Gaussian priors with mean 0 and sd 0.5, and the group-level standard deviation parameters were given positive-truncated Cauchy priors with mean 0 and sd 0.1. Priors for the log gain parameter were the same, except the intercept prior had mean 1 (linear gain estimate of 2.72 in *d*′ units) and sd 1. The posterior distribution represents the model’s beliefs about the parameters given the priors and data. This distribution is summarised above as posterior mean, 95% credible intervals and posterior probabilities for the fixed-effects parameters to be negative (the latter computed via the empirical cumulative distribution of the relevant MCMC samples).

## Acknowledgments

Designed the experiments: TSAW, ASE, CMF, LAG, FAW, MB. Programmed the CNN-model: CMF, LAG. Programmed the experiments: TSAW. Collected the data: CMF, TSAW. Analysed the data: TSAW. Wrote the paper: TSAW, CMF. Revised the paper: ASE, LAG, FAW, MB. Funded by the German Federal Ministry of Education and Research (BMBF) through the Bernstein Computational Neuroscience Program Tübingen (FKZ: 01GQ1002), the German Excellency Initiative through the Centre for Integrative Neuroscience Tübingen (EXC307), and the German Science Foundation (DFG; priority program 1527, BE 3848/2-1 and SFB 1233, Robust Vision: Inference Principles and Neural Mechanisms, TP03). We thank Wiebke Ringels for assistance with data collection, and Heiko Schütt and Corey Ziemba for helpful comments on an earlier draft. TSAW was supported in part by an Alexander von Humboldt Postdoctoral Fellowship. The authors thank the International Max Planck Research School for Intelligent Systems (IMPRS-IS) for supporting Christina Funke.

## Supplementary information for Wallis, Funke et al.

Wallis, Funke, Ecker, Gatys, Wichmann & Bethge

### 1 Freeman and Simoncelli: additional experiments

#### 1.1 Image difficulty

The primary manuscript presented data for 10 “scene-like” and 10 “texture-like” images (Figure S2). The different images within these categories showed more variance in scale factor than did the participants in the experiment (Figure 2G in the primary manuscript, posterior means for each image as diamonds). Here we show the images with the lowest and highest scale factors within each image type category.

In general the results (Figure S3) affirm that our classification into “scene-like” and “texture-like” images is not very precise. The scene-like image with the lowest critical scale factor is one in which a large contour / discontinuity in structure (the border between sky and buildings) passes right through the centre of the image (and thus the fovea). This seems interpretable: participants are very sensitive to any disruption of this prominent contour. Indeed, for this image it does not really make sense to talk about a “critical scale”, because the data show that participants are always very sensitive to the difference between the original image and the syntheses. The critical scale estimate in this case is more a product of the model than of the data, which do not constrain the estimate. On the other hand, the scene-like image with the highest critical scale also contains a prominent contour quite close to the fovea. It is unclear why syntheses for this image are relatively successful at producing indiscriminable samples. Similarly, for the texture-like images, the graffiti image with the lowest critical scale (i.e. for which the model must retain relatively more information to produce indiscriminable samples) is subjectively more “texture-like” than the image with the highest critical scale (landscape image), which contains several long, sharp edges delimiting different structure in the image. Again, if our intuition on the determinant image features is correct, one might expect the opposite pattern of results.

Overall, these results urge caution with respect to the image features that are critical for when the FS model succeeds or fails. While we indeed show average differences between so-called “scene-like” and “texture-like” images, there are individual images for which this interpretation does not hold. Nevertheless, the overall data support the the observation that the “metamers” produced by encoding models that pool texture-like features over fixed pooling regions depend on the image content.

**Figure S1.**
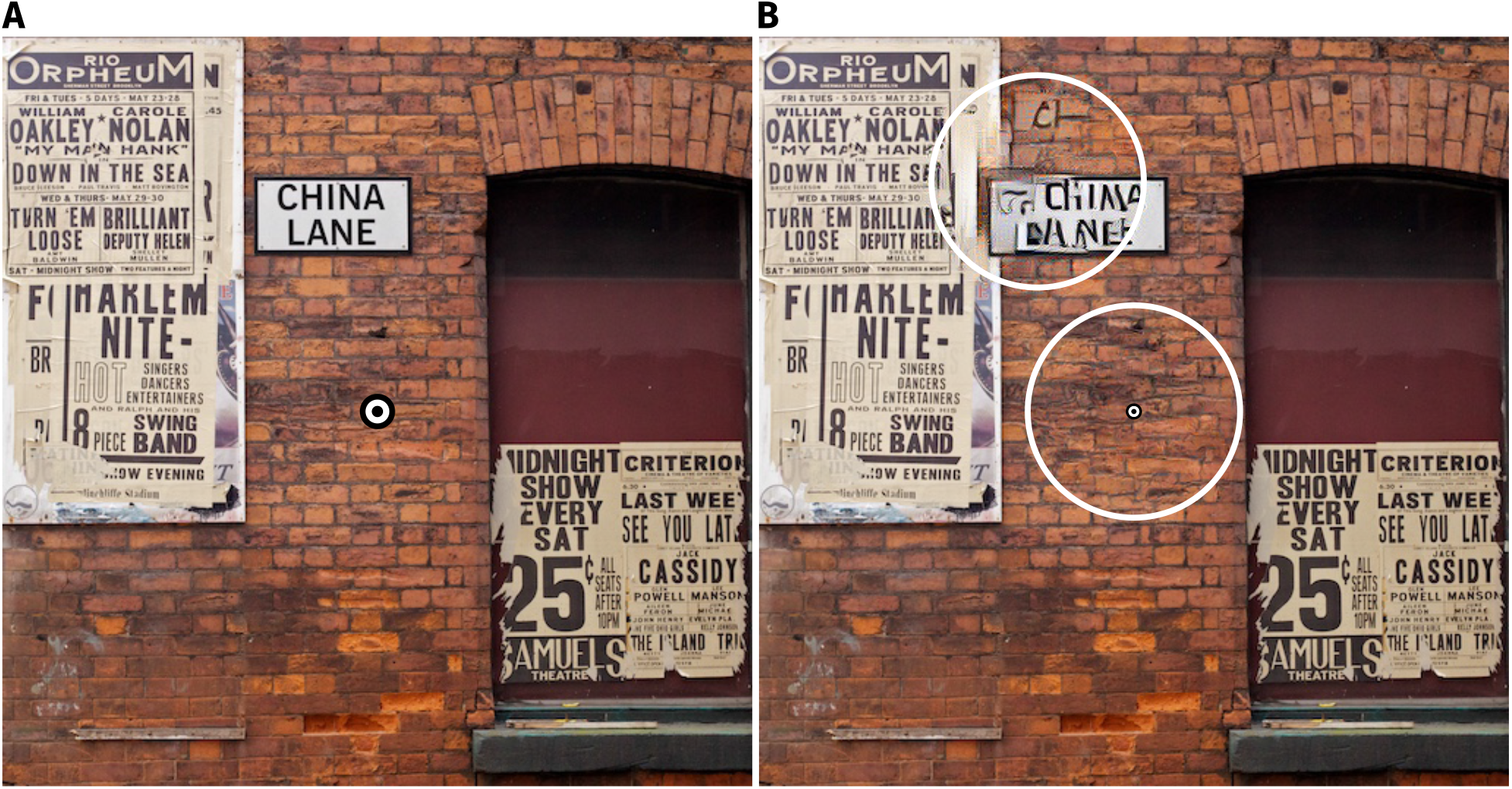
Higher-resolution versions of the images from Figure 1 of the main paper. A: “Geotemporal Anomaly” by Pete Birkinshaw (2010; re-used under a CC-BY 2.0 license). B: The image from A with two circular texture-like distortions (circles enclose distorted area).

**Figure S2.**
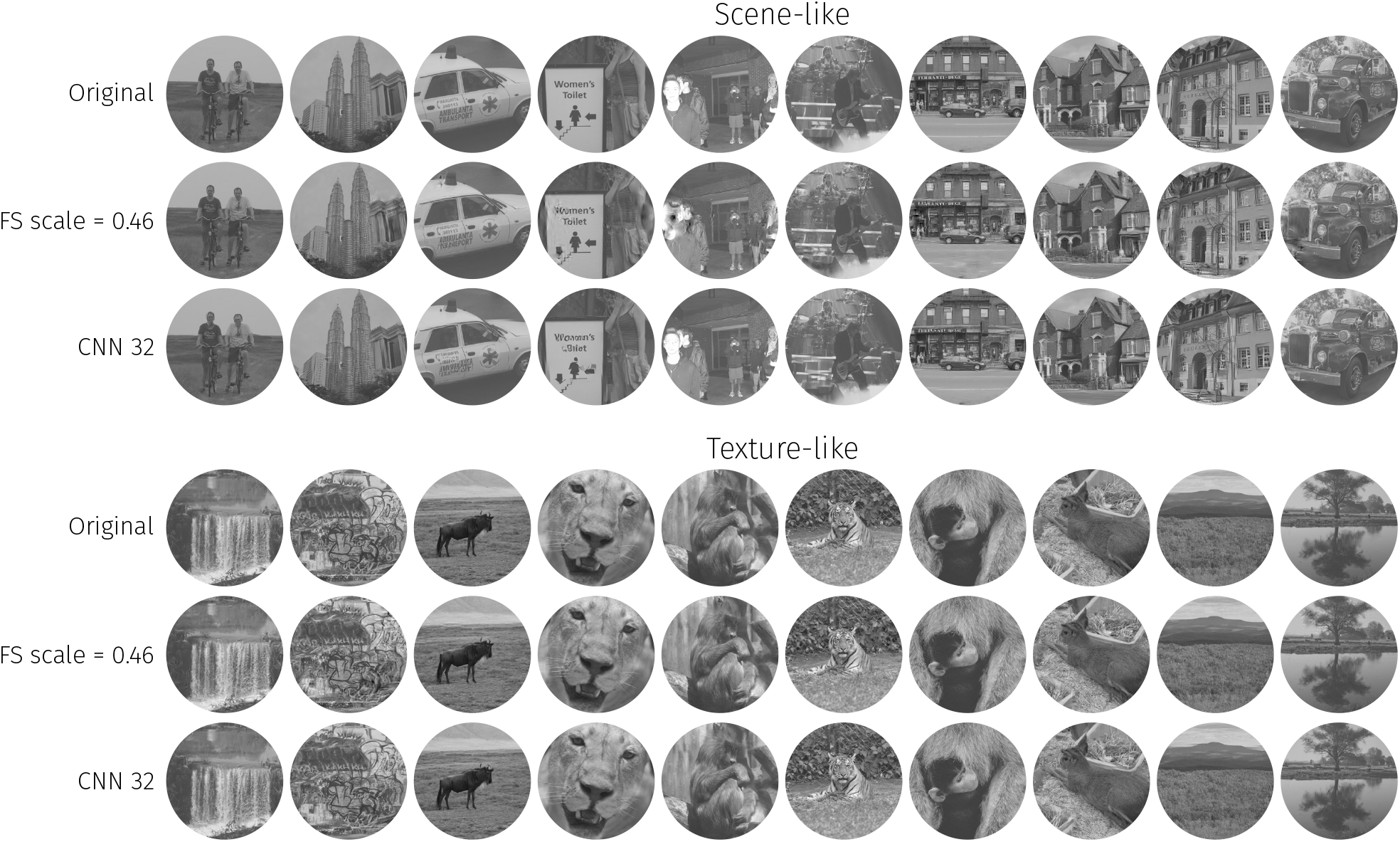
The ten scene-like and ten texture-like images used in our main experiments, along with example syntheses from the FS-0.46 and CNN 32 models (best viewed with zoom).

**Figure S3.**
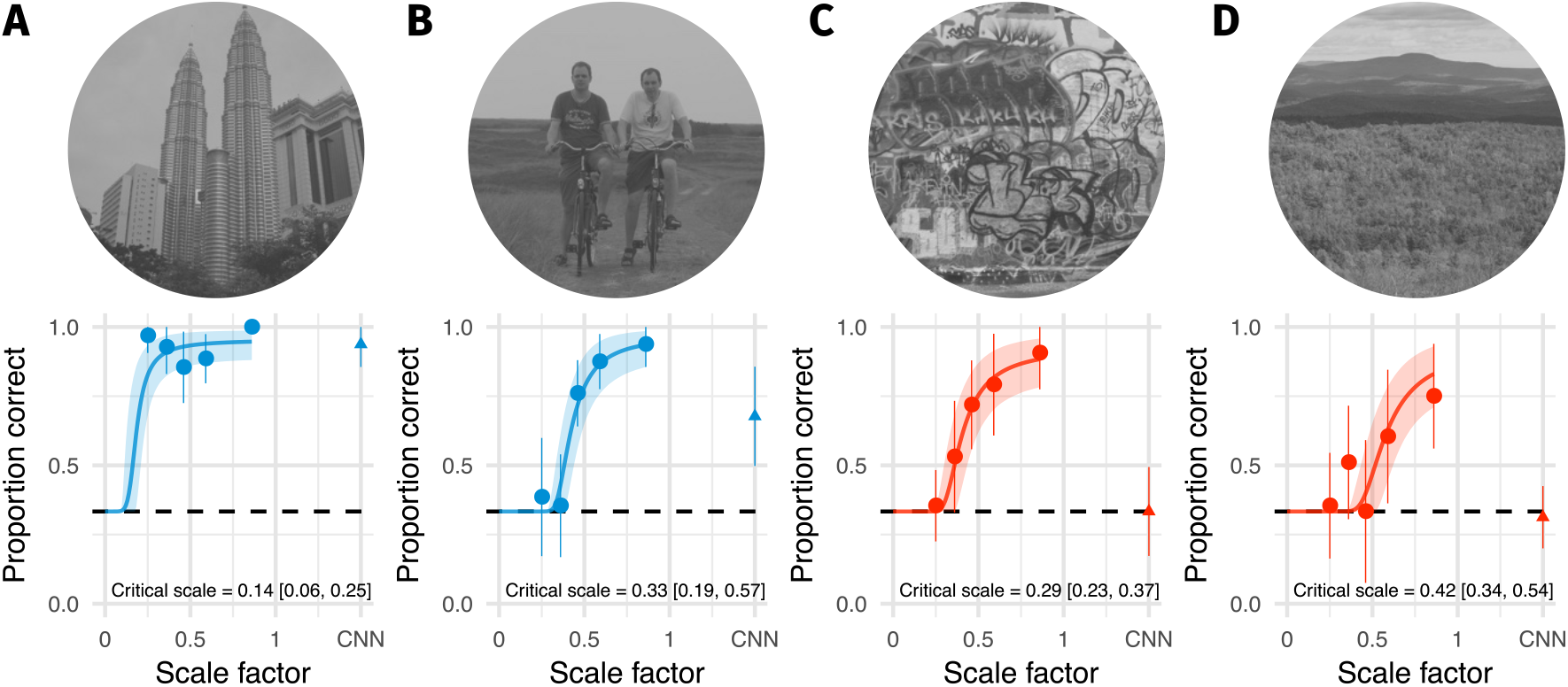
Within the categories of image type, some images have lower critical scales than others (shown for orig vs synth comparison). A: Scene-like image with the lowest critical scale estimate. Points and error bars show grand mean and ±2 SE over participants, solid curve and shaded area show posterior mean and 95% credible intervals from the mixed-effects model. Inline text shows posterior mean and 95% credible interval on the critical scale estimate for this image. B: Scene-like image with the highest critical scale estimate. C: Texture-like image with the lowest critical scale estimate. D: Texture-like image with the highest critical scale estimate.

#### 1.2 Stimulus artifact control

During the course of our testing we noticed that synthesised images generated with the code from http://github.com/freeman-lab/metamers contained an artifact, visible as a wedge in the lower-left quadrant of the synthesised images in which the phases of the surrounding image structure were incorrectly matched (Figure S4A). The angle and extent of the wedge changed with the scale factor, and corresponded to the region where angular pooling regions wrapped from 0–2π (Figure S4B–C). The visibility of the artifact depended on image structure, but was definitely due to the synthesis procedure itself because it also occurred when synthesising matches to a white noise source image (Figure S4D–E). The artifact was not peculiar to our hardware or implementation because it is also visible in the stimuli shown in Deza et al (2017).

Participants in our experiment could have learned to use the artifact to help discriminate images, particularly synthesised images from original images (since only synthesised images contain the artifact). This may have boosted their sensitivity more than might be expected from the model described by Freeman and Simoncelli, leading to the lower critical scales we observed. To control for this, we re-ran the original vs synth condition with the same participants, with the exception that the lower-left quadrant of the image containing the artifact was masked by a grey wedge (in both original and synthesised images) with angular subtense of 60 degrees. We used only the lowest two scale factors from the main experiment, and participants completed this control experiment after the main experiment reported in the paper. We discarded trials for which participants made no response (N = 9) or broke fixation (N = 57), leaving a total of 1014 trials for further analysis. If the high sensitivity at low scale factors we observed above were due to participants using the artifact, then their performance with the masked stimuli should fall to chance for low scale factors.

This is not what we observed: while performance with the wedge was slightly worse (perhaps because a sizable section of the image was masked), the scene-like images remained above chance performance for the lowest two scale factors (Figure S4F). This shows that the low critical scale factors we observed in the main experiment are not due to the wedge artifact.

**Figure S4.**
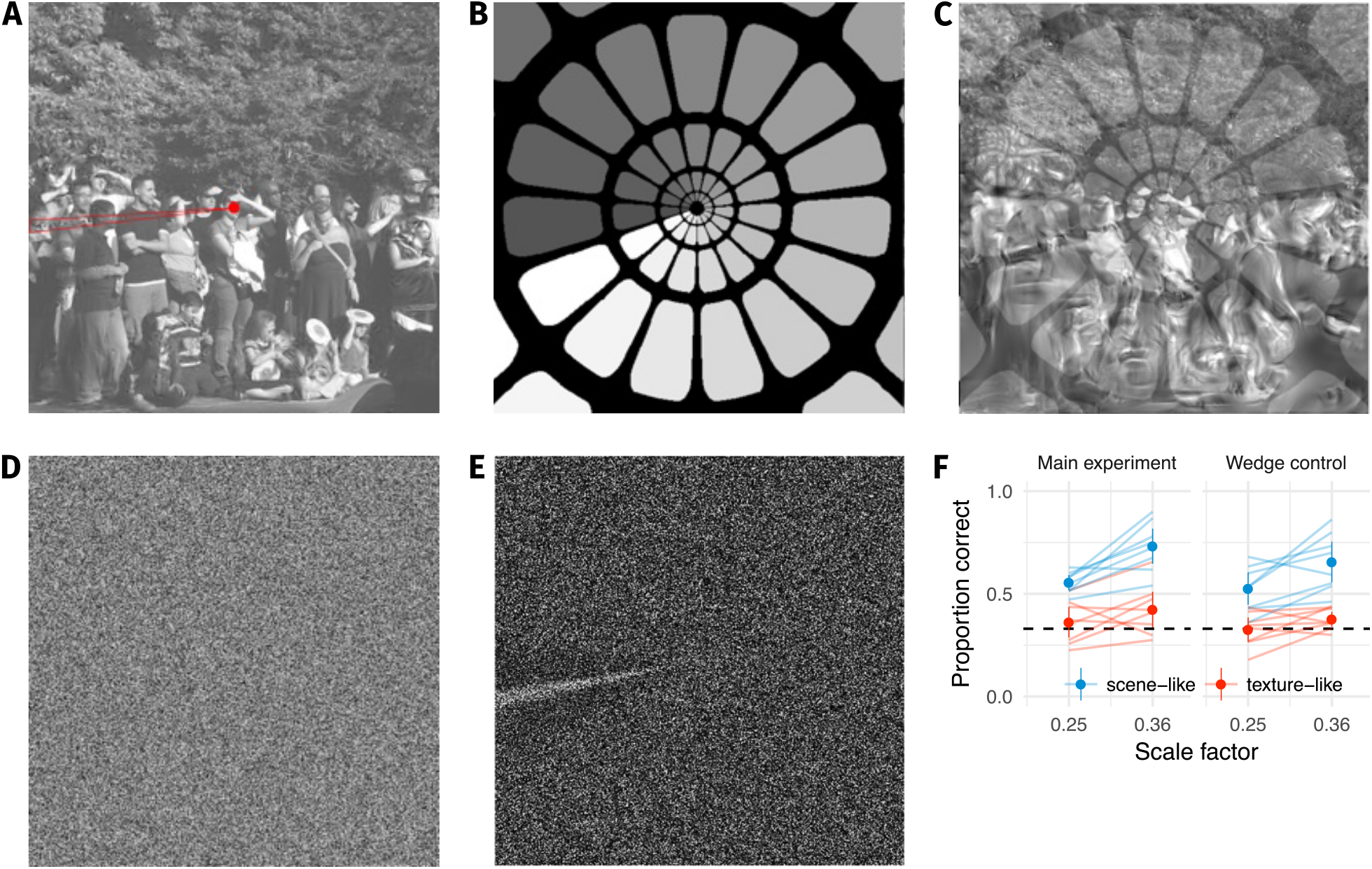
Our results do not depend on an artifact in the synthesis procedure. A: During our pilot testing, we noticed a wedge-like artifact in the synthesis procedure of Freeman and Simoncelli (highlighted in red wedge). The artifact occurred where the angular pooling regions wrapped from 0 to 2π (B: pooling region contours shown with increasing greyscale to wrap point, C: overlayed on scene with artifact). D: The artifact was not driven by image content, because it also occurred when synthesising to match white noise (shown with enhanced contrast in E). If participants’ good performance at small scale factors was due to taking advantage of this wedge, removing it by masking out that image region should drop performance to chance. F: Performance at the two smallest scale factors replotted from the main experiment (left) and with a wedge mask overlayed (right) in the orig vs synth comparison. Points show average (±2SE) over participants; faint lines link individual participant means. Performance remains above chance for the scene-like images, indicating that the low critical scales we observed were not due to the wedge artifact.

#### 1.3 ABX control

Participants in our experiment showed poor performance in the synth vs synth condition even for large scale factors (highest accuracy for a participant at the largest scale of 1.45 was 0.8, average accuracy 0.58), leading to relatively flat psychometric functions (Figure 2F of main manuscript). In contrast, most participants in Freeman and Simoncelli (2011) achieved accuracies above 90% correct for the highest scale factor they test (1.45 as in our experiment). One difference between our experiment and Freeman and Simoncelli (2011) is that they used an ABX task, in which participants saw two images A and B, followed by image X, and had to report whether image X was the same as A or B. Perhaps our oddity task is simply harder: due to greater memory load or the cognitive demands of the comparison, participants in our experiment were unable to perform consistently well.

To assess whether the use of an oddity task lead to our finding of lower critical scales and / or poorer asymptotic performance in the synth vs synth condition, we re-ran our experiment as an ABX task. We used the same timing parameters as in Freeman and Simoncelli. Six participants participated in the experiment, including a research assistant (the same as in the main experiment), four naïve participants and author AE (who only participated in the synth vs synth condition). We discarded trials for which participants made no response (N = 61) or broke fixation (N = 442), leaving a total of 7537 trials for further analysis. The predicted proportion correct in the ABX task was derived from *d*′ using the link function given by Macmillan and Creelman (2005, 229–33) for a differencing model in a roving design:

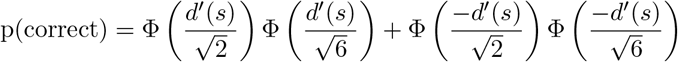

where Φ is the standard cumulative Normal distribution.

As in our main experiment with the oddity task, we find that participants could easily discriminate scene-like syntheses from their original at all scales we could generate (Figure S5). Critical scale factor estimates were similar to those in the main experiment, indicating that the ABX task did not make a large difference to these results. Critical scale estimates were slightly larger, but much more uncertain, in the synth vs synth condition. This uncertainty is largely driven by the even poorer asymptotic performance than in the main experiment. This shows that the results we report in the primary manuscript are not particular to the oddity task.

What explains the discrepancy between asymptotic performance in our experiment vs Freeman and Simoncelli? One possibility is that the participants in Freeman and Simoncelli’s experiment were more familiar with the images shown, and that good asymptotic performance in the synth vs synth condition requires strong familiarity. Freeman and Simoncelli used four original (source) images, and generated three unique synthesised images for each source image at each scale, compared to our 20 source images with four syntheses.

### 2 CNN scene appearance model

Here we describe the CNN scene appearance model presented in the paper in more detail, as well as additional experiemnts concerning this model.

To create a summary statistic model using CNN features, we compute the mean activation in a subset of CNN layers over a number of radial and angular spatial regions (see Figure S6). Increasing the number of pooling regions (reducing the spatial area over which CNN features are pooled) preserves more of the structure of the original image. New images can be synthesised by minimising the difference between the model features for a given input image and a white noise image via an iterative gradient descent procedure (see below). This allows us to synthesise images that are physically different to the original but approximately the same according to the model. We did this for each of four pooling region sizes, named model 4, 8, 16 and 32 respectively after the number of angular pooling regions. These features were matched over three spatial scales, which we found improved the model’s ability to capture long-range correlations.

**Figure S5.**
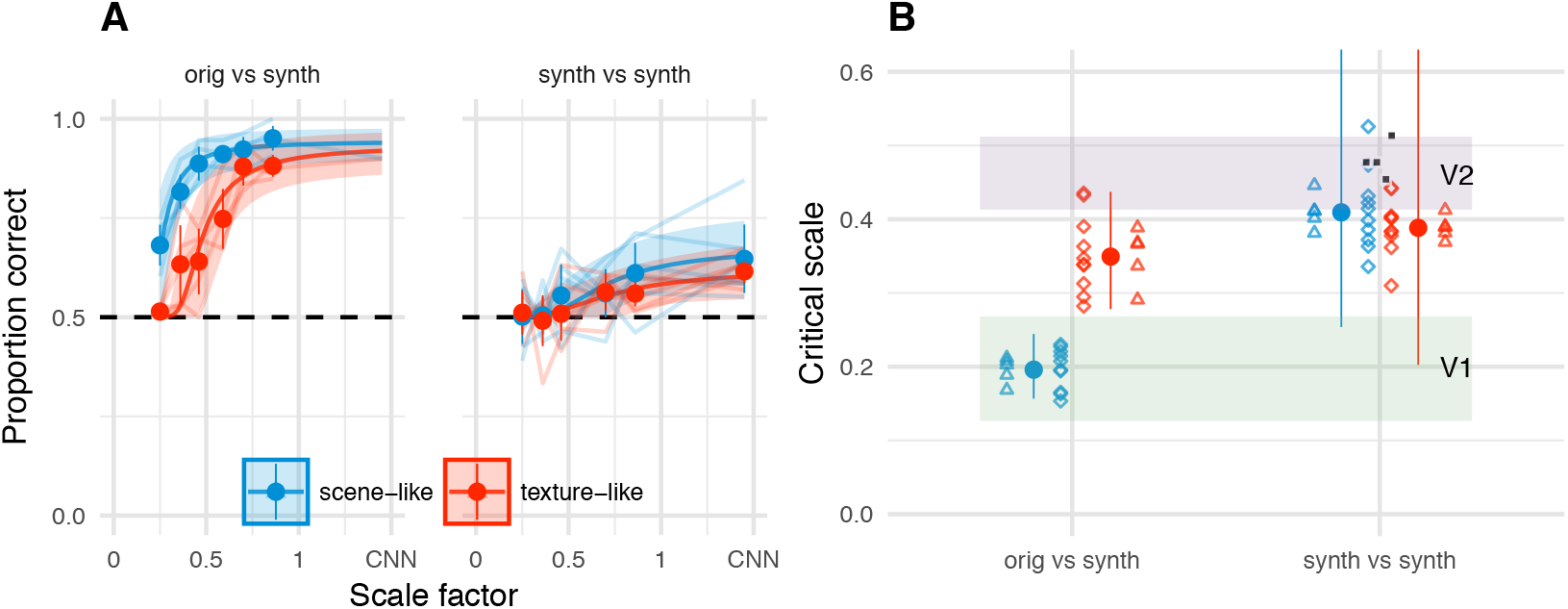
Results from the main paper replicated under an ABX task. A: Performance in the ABX task as a function of scale factor. Points show grand mean ±2 SE over participants; faint lines link individual participant performance levels. Solid curves and shaded regions show the fit of a nonlinear mixed-effects model estimating the critical scale and gain. B: When comparing original and synthesised images, estimated critical scales (scale at which performance rises above chance) are lower for scene-like than for texture-like images. Points with error bars show population mean and 95% credible intervals. Triangles show posterior means for participants; diamonds show posterior means for images. Black squares show critical scale estimates of the four participants from Freeman & Simoncelli reproduced from that paper (x-position jittered to reduce overplotting); shaded regions denote the receptive field scaling of V1 and V2 estimated by Freeman & Simoncelli.

In Experiment 1, we tested the discriminability of syntheses generated from the four pooling models in a set of 400 images that were novel to the participants. Experiment 2 examines the effect of image familiarity by repeatedly presenting a small number of source images. Experiment 3 tested the effect of cueing spatial attention on performance. Experiment 4 tested sensitivity to local texture distortions.

#### 2.1 CNN methods

##### 2.1.1 Radial and angular pooling

In the texture synthesis approach of Gatys et al (2015), spatial information is removed from the raw CNN activations by computing summary statistics (the Gram matrices of correlations between feature maps) over the whole image. In the ‘foveated’ pooling model we present here, we compute and match the mean of the feature maps (i.e. not the full Gram matrices) over local image regions by dividing the image into a number of radial and angular pooling regions (Figure S6). The radius defining the border between each radial pooling region is based on a given number of angular regions *N_θ_* (which divide the circle evenly) and given by

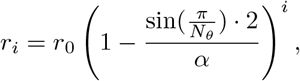

where *r_i_* is the radius of each region *i*, *r_0_* is the outermost radius (set to be half the image size), and – is the ratio between the radial and angular difference. Radial regions were created for all *i* for which *r_i_* Ø 64~px, corresponding to the preserved central region of the image (see below). We set α = 4 because at this ratio *N_θ_* ≈ *N_e_* (where *N_e_* is the number of radial regions) for most *N_θ_*. The value of *N_θ_* corresponds to the model name used in the paper (e.g. ‘CNN 4’ uses *N_θ_* = 4).

We now apply these pooling regions to the activations of the VGG-19 deep CNN (Simonyan and Zisserman 2015). For a subset of VGG-19 layers (conv1_1, conv2_1, conv3_1, conv4_1, conv5_1) we compute the mean activation for each feature map j in each layer l within each (radial or angular) pooling region p as

**Figure S6.**
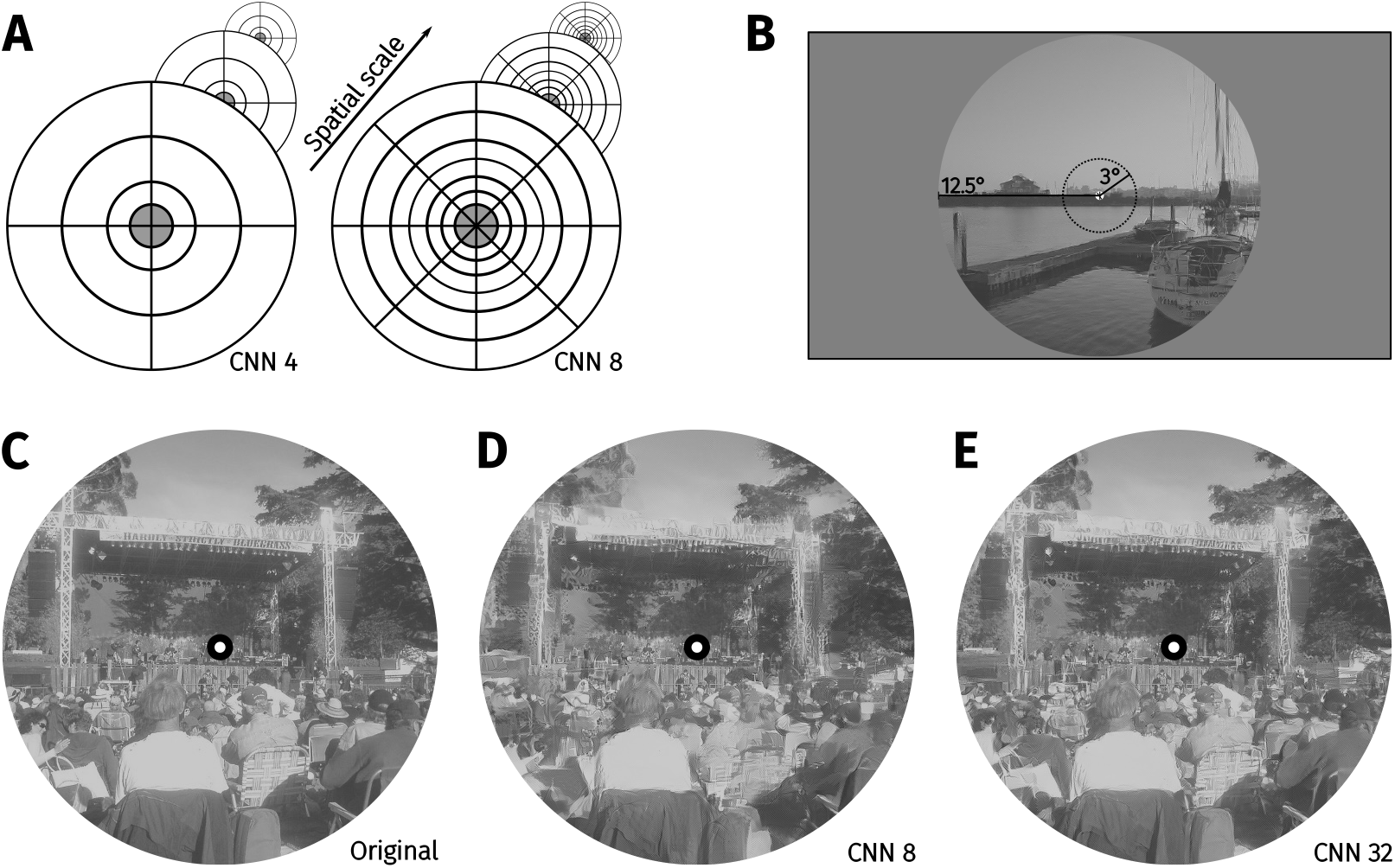
Methods for the CNN scene appearance model. A: The average activations in a subset of CNN feature maps were computed over non-overlapping radial and angular pooling regions that increase in area away from the image centre (not to scale), for three spatial scales. Increasing the number of pooling regions (CNN 4 and CNN 8 shown in this example) increases the fidelity of matching to the original image, restricting the range over which distortions can occur. Higher-layer CNN receptive fields overlap the pooling regions, ensuring smooth transitions between regions. The central 3° of the image (grey fill) is fixed to be the original. B: The image radius subtended 12.5°C: An original image from the MIT1003 dataset. D: Synthesised image matched to the image from C by the CNN 8 pooling model. E: Synthesised image matched to the image from E by the CNN 32 pooling model. Fixating the central bullseye, it should be apparent that the CNN 32 model preserves more information than the CNN 8 model, but that the periphery is nevertheless significantly distorted relative to the original.

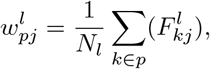

where is *N_l_* the size of the feature map of layer l in pixels and k is the (vectorised) spatial position in feature map 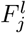. The set of all 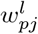 provides parameters that specify the foveated image at a given scale. Note that while the radial and angular pooling region responses are computed separately, because they are added together to the loss function during optimisation (see below) they effectively overlap (as depicted in Figure S6). In addition, while the borders of the pooling regions are hard-edged (i.e. pooling regions are non-overlapping), the receptive fields of CNN units (area of pixels in the image that can activate a given unit in a feature map) can span multiple pooling regions. This means that the model parameters of a given pooling region will depend on image structure lying outside the pooling region (particularly for feature maps in the higher network layers). This encourages smooth transitions between pooling regions in the synthesised images.

##### 2.1.2 Multiscale model

In the VGG-19 network, receptive fields of the units are squares of a certain size, and this size is independent of the input size of the image. That is, given a hypothetical receptive field centred in the image of size 128~px square, the unit will be sensitive to one quarter of the image for input size 512~px but half the image for input size 256. Therefore, the same unit in the network can receive image structure at a different scale by varying the input image size, and in the synthesis process the low (high) frequency content can be reproduced with greater fidelity by using a small (large) input size.

We leverage this relationship to better capture long-range correlations in image structure (caused by e.g. edges that extend across large parts of the image) by computing and matching the model statistics over three spatial scales. This is not a controversial idea: for example, the model of Freeman and Simoncelli (2011) also computes features in a multiscale framework. How many scales is sufficient?

We evaluated the degree to which the number and combination of scales affected appearance in a psychophysical experiment on authors TW and CF. We matched 100 unique images using seven different models: four single-scale models corresponding to input sizes of 64, 128, 256 and 512 pixels, and three multiscale models in which features were matched at multiple scales ([256, 512], [128, 256, 512] and [64, 128, 256, 512]). The foveated pooling regions corresponded to the CNN 32 model. Output images were upsampled to the final display resolution as appropriate. We discarded trials for which participants made no response (N = 2) or broke fixation (N = 5), leaving a total of 1393 trials for further analysis.

Figure S7 shows that participants are sensitive to the difference between model syntheses and original images when features are matched at only a single scale. However, using two or three scales appears to be sufficient to match appearance on average. As a compromise between fidelity and computational tractability, we therefore used three scales for all other experiments on the CNN appearance model. The final model used three networks consisting of the same radial and angular regions described above, computed over sizes 128, 256 and 512~px square. The final model representation W therefore consists of the pooled feature map activations over three scales: 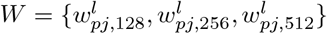.

##### 2.1.3 Gradient descent

As in Gatys et al (2015), synthesised images are generated using iterative gradient descent, in which the mean-squared distance between the averaged feature maps of the original image and the synthesis is minimized. If *T* and *W* are the model representations for the synthesis and the original image respectively, then the loss for each layer is given by

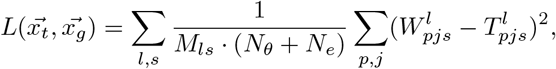

**Figure S7.**
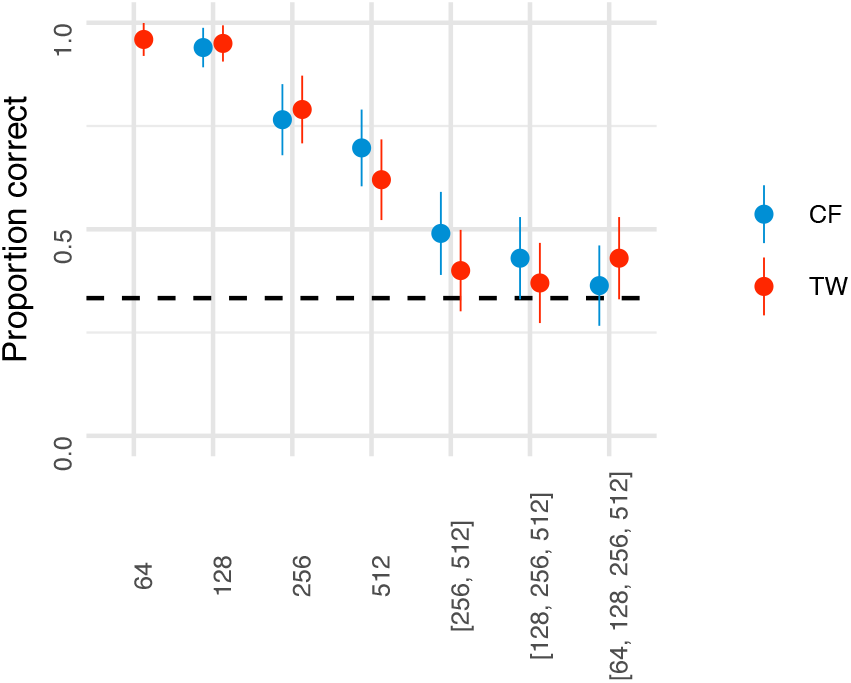
Performance for discriminating model syntheses and original scenes for single- and multi-scale models (all with pooling regions corresponding to CNN 32) for participants CF and TW. Points show participant means (error bars show ±2 SEM), dashed line shows chance performance. The multiscale model with three scales produces close-to-chance performance.

where 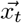 and 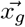 are the vectorised pixels of the original and new image respectively, *M_l,s_* is the number of feature maps for layer *l* in scale s. A circular area in the middle of the image (radius 64~px) is preserved to be the original image. Tiling pooling regions even for the centre of the image created reasonable syntheses but is prohibitively costly in generation time. To preserve the pixels in the circular area, the initialisation image of the gradient descent is identical to the original image. Outside the central area the gradient descent is initialised with Gaussian noise. The gradient descent used the L-BFGS optimiser (scipy implementation, Jones, Oliphant, and Peterson 2001) for 1000 iterations.

#### 2.2 Experiment 1: Discriminability of CNN model syntheses for 400 unique images

This experiment measured whether any of the variants of the CNN scene appearance model could synthesise images that humans could not discriminate from their natural source images, and if so, identify the simplest variant of this model producing metamers. We chose a set of 400 images and had participants discriminate original and model-generated images in a temporal oddity paradigm.

##### 2.2.1 Methods

The methods for this and the following psychophysical experiments were the same as in the main paper unless otherwise noted.

###### 2.2.1.1 Participants

Thirteen participants participated in this experiment. Of these, ten participants were recruited via online advertisements and paid 15 Euro for a 1.5 hour testing session; the other three participants were authors AE, TW and CF. One session comprised one experiment using unique images (35 mins) followed by and one of repeated images (see below; 25 mins). All participants signed a consent form prior to participating. Participants reported normal or corrected-to-normal visual acuity. All procedures conformed to Standard 8 of the American Psychological Association’s “Ethical Principles of Psychologists and Code of Conduct” (2010).

###### 2.2.1.2 Stimuli

We used 400 images (two additional images for authors, see below) from the MIT 1003 database (Judd, Durand, and Torralba 2012; Judd et al. 2009). One of the participants (TW) was familiar with the images in this database due to previous experiments. New images were synthesised using the multiscaled (512 px, 256 px, 128 px) foveated model described above, for four pooling region complexities (4, 8, 16 and 32). An image was synthesised for each of the 400 original images from each model (giving a total stimulus set including originals of 2000).

###### 2.2.1.3 Procedure

Participants viewed the display from 60 cm; at this distance, pixels subtended approximately 0.024 degrees on average (approximately 41 pixels per degree of visual angle) – note that this is slightly further away than the experiment reported in the primary paper (changed to match the angular subtense used by Freeman and Simoncelli). Images therefore subtended ≈ 12.5° at the eye. As in the main paper, the stimuli were presented for 200 ms, with an inter-stimulus interval of 1000 ms, followed by a 1200 ms response window. Feedback was provided by a 100 ms change in fixation cross brightness. Gaze position was recorded during the trial. If the participant moved the eye more than 1.5 degrees away from the fixation spot, feedback signifying a fixation break appeared for 200~ms after the response feedback. Prior to the next trial, the state of the participant’s eye position was monitored for 50 ms; if the eye position was reported as more than 1.5 degrees away from the fixation spot a recalibration was triggered. The inter-trial interval was 400 ms.

Each unique image was assigned to one of the four models for each participant (counterbalanced). That is, a given image might be paired with a CNN 4 synthesis for one participant and a CNN 8 synthesis for another. Showing each unique image only once ensures that the participants cannot become familiar with the images. For authors, images were divided into only CNN 8, CNN 16 and CNN 32 (making 134 images for each model and 402 trials in total for these participants). To ensure that the task was not too hard for naïve participants we added the easier CNN 4 model (making 100 images for each model version and 400 trials in total). The experiment was divided into six blocks consisting of 67 trials (65 trials for the last block). After each block a break screen was presented telling the participant their mean performance on the previous trials. During the breaks the participants were free to leave the testing room to take a break and to rest their eyes. At the beginning of each block the eyetracker was recalibrated. Naïve participants were trained to do the task, first using a slower practice of 6 trials and second a correct-speed practice of 30 trials (using five images not part of the stimulus set for the main experiment).

###### 2.2.1.4 Data analysis

We discarded trials for which participants made no response (N = 81) or broke fixation (N = 440), leaving a total of 4685 trials for further analysis.

Performance at each level of CNN model complexity was quantified using a logistic mixed-effects model. Correct responses were assumed to arise from a fixed effect factor of CNN model (with four levels) plus the random effects of participant and image. The model (in lme4-style notation) was

`correct ~ model + (model | subj) + (model | im_code)`

with `family = Bernoulli(“logit”)`, and using `contr.sdif` coding for the CNN model factor (Venables and Ripley 2002).

The posterior distribution over model parameters was estimated using weakly-informative priors, which provide scale information about the setting of the model but do not bias effect estimates above or below zero. Specifically, fixed effect coefficients were given Cauchy priors with mean zero and SD 1, random effect standard deviations were given bounded Cauchy priors with mean 0.2 (indicating that we expect some variance between the random effect levels) and SD 1, with a lower-bound of 0 (variances cannot be negative), and correlation matrices were given LKJ(2) priors, reflecting a weak bias against strong correlations (Stan Development Team 2015). The model posterior was estimated using MCMC implemented in the Stan language (version 2.16.2, Stan Development Team 2017; Hoffman and Gelman 2014), with the model wrapper package brms (version 1.10.2, Bürkner 2017) in the R statistical environment. We computed four chains of 15,000 steps, of which the first 5000 steps were used to tune the sampler; to save disk space we only saved every 5th sample.

**Figure S8.**
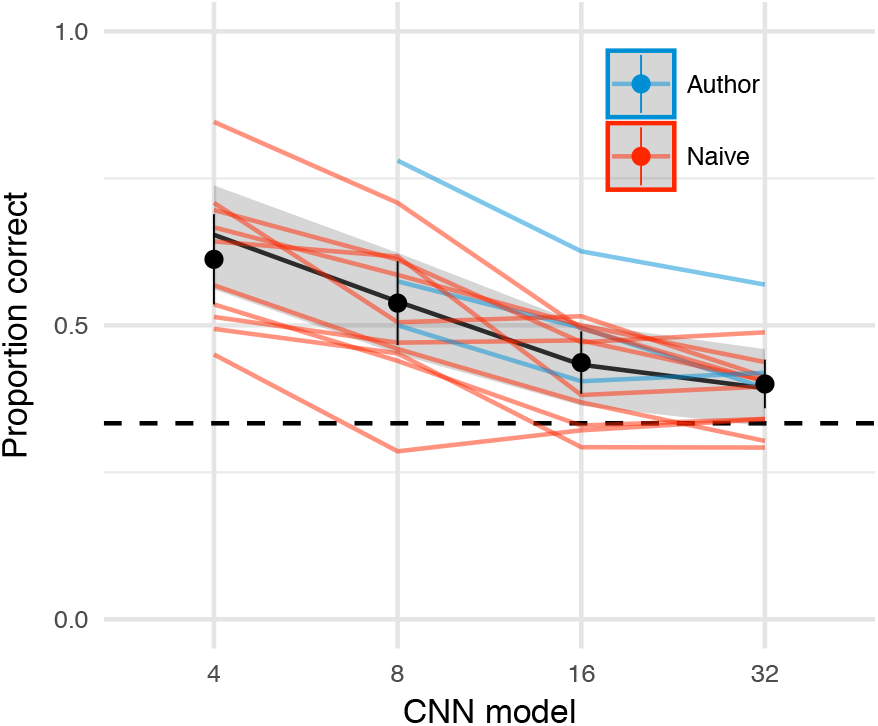
The CNN model comes close to matching appearance on average. Oddity performance as a function of the CNN image model. Points show mean over participants (error bars ±2 SEM), coloured lines link the mean performance of each participant for each pooling model. For most participants, performance falls to approximately chance (dashed horizontal line) for the CNN 32 model. Black line and shaded regions show the mean and 95% credible intervals on the population mean derived from a mixed-effects model.

##### 2.2.2 Results and discussion

The CNN 32 model came close to matching appearance on average for a set of 400 images. Discrimination performance for ten naïve participants and three authors is shown in Figure S8 (lines link individual participant means, based on at least 64 trials, median 94). All participants achieve above-chance performance for the simplest model (CNN 4), indicating that they understood and could perform the task. Performance deteriorates as models match the structure of the original image more precisely.

To quantify the data, we estimated the posterior distribution of a logistic mixed-effects model with a population-level (fixed-effect) factor of CNN model, whose effect was nested within participants and image (i.e. random effects of participant and image). Regression coefficients coded the difference between successive CNN models, expressed using sequential difference coding from the MASS package (Venables and Ripley 2002), and are presented below as the values of the linear predictor (corresponding to log odds in a logistic model). Mean performance had a greater than 0.99 posterior probability of being lower for CNN 8 than CNN 4 (-0.48, 95% CI [-0.74, -0.23], p(*β* < 0) > 0.999), and for CNN 16 being lower than CNN 8 (-0.44, 95% CI [-0.68, -0.18], p(*β* < 0) = 0.999); whereas the difference between the 16 and 32 models was somewhat smaller (-0.17, 95% CI [-0.37, 0.03], p(*β* < 0) = 0.951). Most participants performed close to chance for the CNN 32 model (excluding authors, the population mean estimate had a 0.88 probability of being greater than chance; including authors this value was 0.96). Therefore, the model is capable of synthesising images that are indiscriminable from a large set of arbitrary scenes in our experimental conditions, on average, for naïve participants. However, one participant (author AE) performs noticably better than the others, even for the CNN 32 model. AE had substantial experience with the type of distortions produced by the model but had never seen this set of original images before. Therefore, the images produced by the model are not true metamers, because they do not encapsulate the limits of visible structure for all humans.

#### 2.3 Experiment 2: Image familiarity and learning tested by repeated presentation

It is plausible that familiarity with the images played a role in the results above. That is, the finding that images become difficult on average to discriminate with the CNN 32 model may depend in part on participants having never seen the images before. To investigate the role that familiarity with the source images might play, the same participants as in the experiment above performed a second experiment in which five of the original images from the first experiment were presented 60 times, using 15 unique syntheses per image generated with the CNN 32 model (Figure S9A).

##### 2.3.1 Methods

###### 2.3.1.1 Participants

The same thirteen participants participated as in Experiment 1.

###### 2.3.1.2 Stimuli

We selected five images from the set of 400 and generated 15 new syntheses for each of these images from the CNN 32 model (yielding a stimulus set of 80 images).

###### 2.3.1.3 Procedure

Each pairing of unique image (5) and synthesis (15) was shown in one block of 75 trials (pseudo-random order with the restriction that trials from the same source image could never follow one another). Participants performed four such blocks, yielding 300 trials in total (60 repetitions of each original image).

###### 2.3.1.4 Data analysis

We discarded trials for which participants made no response (N = 63) or broke fixation (N = 294), leaving a total of 3543 trials for further analysis. Model fitting was as for Experiment 1 above, except that the final posterior was based on four chains of 16,000 steps, of which the first 8000 steps were used to tune the sampler; to save disk space we only saved every 4th sample.

The intercept-only model (assuming only random effects variation but no learning) was specified as

`correct ~ 1 + (1 | subj) + (1 | im_name)`

and the learning model was specified as

`correct ~ session + (session | subj) + (session | im_name)`

We compare models using an information criterion (LOOIC, Vehtari et al (2016); see also (Gelman, Hwang, and Vehtari 2014; McElreath 2016)) that estimates of out-of-sample prediction error on the deviance scale.

##### 2.3.2 Results and discussion

While some images (e.g. House) could be discriminated quite well by most participants (Figure S9B), others (e.g. Graffiti) were almost indiscriminable from the model image for all participants (posterior probability that the population mean was above chance performance was 0.61 for Graffiti, 0.93 for Market, and greater than 0.99 for all other images). This image dependence shows that even the CNN 32 model is insufficient to produce metamers for arbitrary scenes.

Furthermore, there was little evidence that participants learned over the course of sessions (Figure S9C). The population-level linear slope of session number was 0.03, 95% CI [-0.1, 0.15], p(*β* < 0) = 0.326, and the LOOIC comparison between the intercept-only model and the model containing a learning term indicated equivocal evidence if random-effects variance was included (LOOIC difference 3.3 in favour of the learning model, SE = 6.1) but strongly favoured the intercept model if only fixed-effects were considered (LOOIC difference -23.3 in favour of the intercept model, SE = 1.7). The two images with the most evidence for learning were Children (median slope 0.04, 95% CI [-0.08, 0.17], p(*β* < 0) = 0.247) and Sailboat (0.04, 95% CI [-0.08, 0.17], p(*β* < 0) = 0.269). Two authors showed some evidence of learning: AE (0.17, 95% CI [-0.03, 0.37], p(*β* < 0) = 0.047), and CF (0.22, 95% CI [0.03, 0.44], p(*β* < 0) = 0.008). Overall, these results show that repeated image exposures with response feedback did not noticably improve performance.

**Figure S9.**
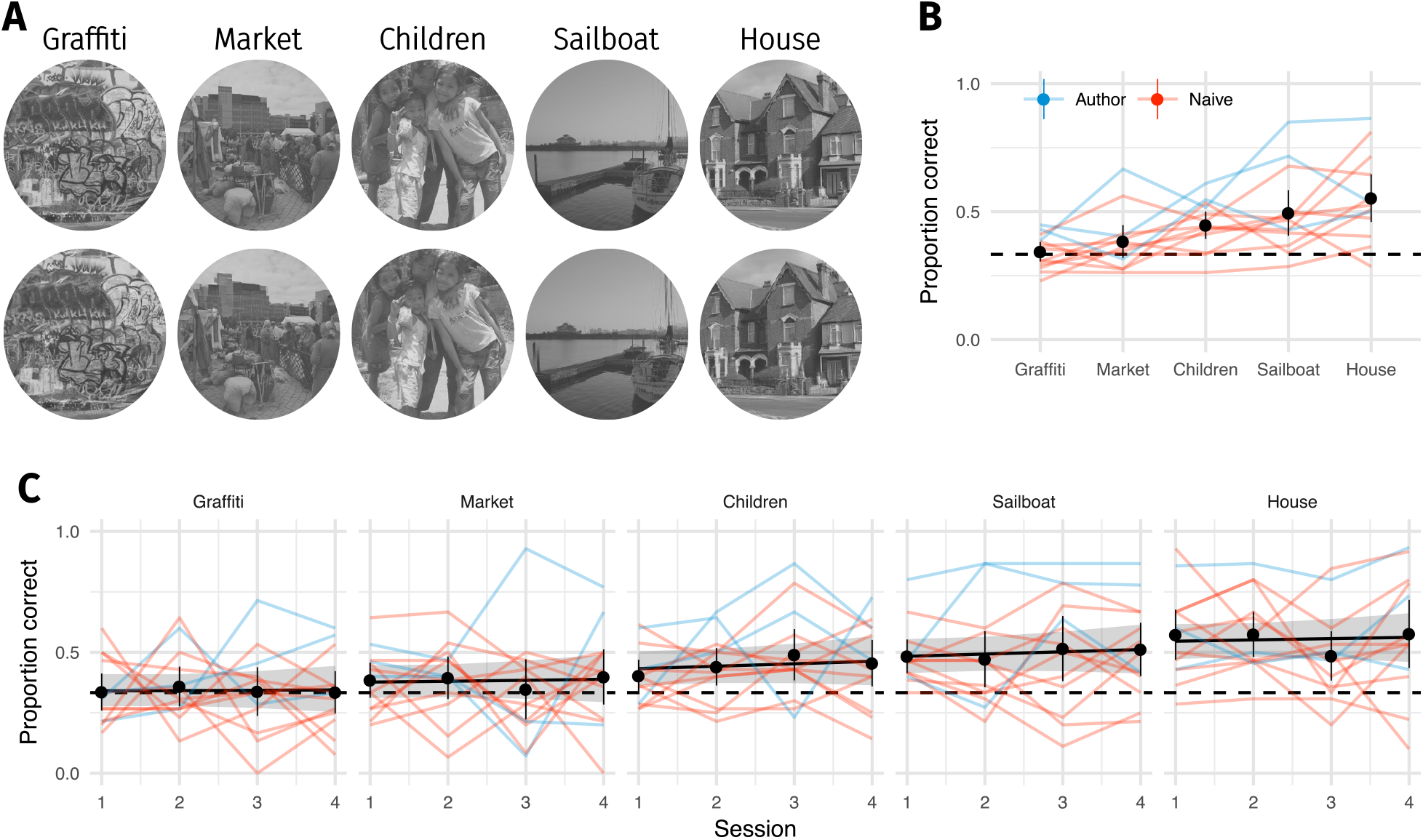
**A:** Five original images (top) were repeated 60 times (interleaved over 4 blocks), and observers discriminated them from CNN 32 model syntheses (bottom). **B:** Proportion of correct responses for each image from **A**. Some images are easier than others, even for the CNN 32 model. **C:** Performance as a function of each 75-trial session reveals little evidence that performance improves with repeated exposure. Points show grand mean (error bars show bootstrapped 95% confidence intervals), lines link the mean performance of each observer for each pooling model (based on at least 5 trials; median 14). Black line and shaded region shows the posterior mean and 95% credible intervals of a logistic mixed-effects model predicting the population mean performance for each image.

#### 2.4 Experiment 3: Spatial cueing of attention

The experiment presented in the primary paper showed that the discriminability of model syntheses depended on the source images, with scene-like images being easier to discriminate from model syntheses than texture-like images for a given image model. This finding was replicated in an ABX paradigm (above) and the general finding of source-image-dependence was corroborated by the data with repeated images (Figure S8). One possible reason for this image-dependence could be that participants found it easier to know where to attend in some images than in others, creating an image-dependence not due to the summary statistics per se. Relatedly, Cohen and co-authors (2016) suggest that the limits imposed by an ensemble statistic representation can be mitigated by the deployment of spatial attention to areas of interest. Can the discriminability of images generated by our model be influenced by focused spatial attention?

To probe this possibility we cued participants to a spatial region of the scene before the trial commenced. We computed the mean squared error (MSE) between the original and synthesised images within 12 partially-overlapping wedge-like regions subtending 60°. We computed MSE in both the pixel space (representing the physical difference between the two images) and in the feature space of the fifth convolutional layer (conv5_1) of the VGG-19 network, with the hypothesis that this might represent more perceptually relevant information, and thus be a more informative cue.

We pre-registered the following hypotheses for this experiment (available at http://dx.doi.org/10.17605/OSF.IO/MBGSQ; click on “View Registration Form”). For the overall effect of cueing (the primary outcome of interest), we hypothesised that

- performance in the Valid:Conv5 condition would be higher than the Uncued condition and
- performance in the Invalid condition would be lower than the Uncued condition

These findings would be consistent with the account that spatial attention can be used to overcome ensemble statistics in the periphery, providing that it is directed to an informative location. This outcome also assumes that our positive cues (Conv5 and Pixels) identify informative locations.

Alternative possibilities are

- if focussed spatial attention cannot influence the “resolution” of the periphery in this task, then performance in the Valid:Conv5 and Invalid conditions will be equal to the Uncued condition.
- if observers use a global signal (“gist”) to perform the task, performance in the Uncued condition would be higher than the Valid:Conv5 and Invalid conditions. That is, directing spatial attention interferes with a gist cue.

Our secondary hypothesis concerns the difference between Valid:Conv5 and Valid:Pixel cues. A previous analysis at the image level (see below) found that conv5 predicted image difficultly slightly better than the pixel space. We therefore predicted that Valid spatial cues based on Conv5 features (Valid:Conv5) should be more effective cues, evoking higher performance, than Valid:Pixel cues.

##### 2.4.1 Methods

###### 2.4.1.1 Participants

**Figure S10.**
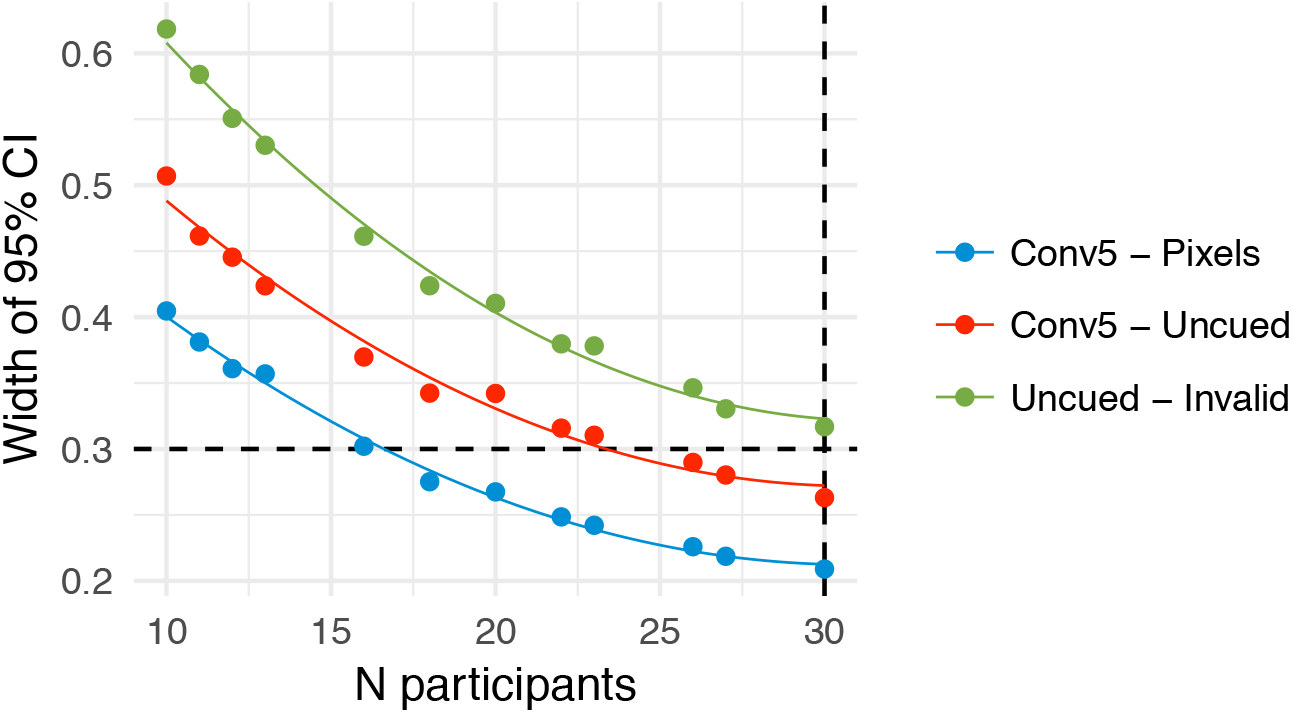
Parameter precision as a function of number of participants. **A:** Width of the 95% credible interval on three model parameters as a function of the number of participants tested. Points show model fit runs (the model was not re-estimated after every participant due to computation time required). We aimed to achieve a width of 0.3 (dashed horizontal line) on the linear predictor scale, or stop after 30 participants. The Uncued - Invalid parameter failed to reach the desired precision after 30 participants. Lines show fits of a quadratic polynomial as a visual guide.

We pre-registered (http://dx.doi.org/10.17605/OSF.IO/MBGSQ) the following data collection plan with a stopping rule that depended on the precision (Kruschke 2015). Specifically, we collected data from a minimum of 10 and a maximum of 30 participants, planning to stop in the intermediate range if the 95% credible intervals for the two parameters of interest (population fixed-effect difference between Valid and Uncued, and population fixed-effect difference between Invalid and Uncued) spanned a width of 0.3 or less on the linear predictor scale.

This value was determined as 75% of the width of our “Region of Practical Equivalence” to zero effect (ROPE), pre-registered as [-0.2, 0.2] on the linear predictor scale (this corresponds to odds ratios of [0.82, 1.22]). We deemed any difference smaller than this value as being too small to be practically important. As an example, if the performance in one condition is 0.5, then an increase of 0.2 in the linear predictor corresponds to a performance of 0.55. The target for precision was then determined as 75% of the ROPE width, in order to give a reasonable chance for the estimate to lie within the ROPE (Kruschke 2015).

We tested these conditions by fitting the data model (see below) after every participant after the 10th, stopping if the above conditions were met. However, as shown in Figure S10, this precision was not met with our maximum of 30 participants, and so we ceased data collection at 30, deeming further data collection beyond our resources for the experiment. Thus our data should be interpreted with the caveat that the desired precision was not reached (though we got close).

An additional five participants were recruited but showed insufficient eyetracking accuracy or training performance (criteria pre-registered). Of the 30, three were lab members unfamiliar with the purpose of the study, the other 27 were recruited online; all were paid 15 Euro for the 1.5 hour testing session. Of these, three participants did not complete the full session due to late arrival, and eyetracking calibration failed in the second last trial block for an additional participant.

###### 2.4.1.2 Stimuli

This experiment used the same 400 source images and CNN 8 model syntheses as Experiment 1.

###### 2.4.1.3 Procedure

The procedure for this experiment was as in Experiment 1 with the following exceptions. The same 400 original images were used as in Experiment 1, all with syntheses from the CNN 8 model. A trial began with the presentation of a bright wedge (60 degree angle, Weber contrast 0.25) or circle (radius 2 dva) for 400~ms, indicating a spatial cue (85% of trials) or Uncued trial (15%) respectively (Figure S11A). A blank screen with fixation spot was presented for 800 ms before the oddity paradigm proceeded as above. On spatial cue trials, participants were cued to the wedge region containing either the largest pixel MSE between the original and synthesised images (35% of all trials), the largest conv5 MSE (35%), or the *smallest* pixel MSE (an invalid cue, shown on 15% of all trials). Thus, 70% of all trials were valid cues, encouraging participants to make use of the cues rather than learning to ignore them. Participants were also instructed to attend to the cued region on trials where a wedge was shown. For Uncued trials they were instructed to attend globally over the image. Cueing conditions were interleaved and randomly assigned to each unique image for each participant. The experiment was divided into eight blocks of 50 trials. Before the experiment we introduced participants to the task and fixation control with repeated practice sessions of 30 trials (using 30 images not used in the main experiment and with the CNN 4 model syntheses). Participants saw at least 60 and up to 150 practice trials, until they were able to get at least 50% correct and with 20% or fewer trials containing broken fixations or blinks.

###### 2.4.1.4 Data analysis

We discarded trials for which participants made no response (N = 141) or broke fixation (N = 1398), leaving a total of 10261 trials for further analysis.

This analysis plan was pre-registered and is available at http://dx.doi.org/10.17605/OSF.IO/MBGSQ (click on “view registration form”). We seek to estimate three performance differences:

1. The difference between Invalid and Uncued
2. The difference between Valid:Conv5 and Uncued
3. The difference between Valid:Conv5 and Valid:Pixels

The model formula (in lme4-style formula notation) is

`correct ~ cue + (cue | subj) + (cue | im_code)`

with `family = Bernoulli(“logit”)`. The “cue” factor uses custom contrast coding (design matrix) to test the hypotheses of interest. Specifically, the design matrix for the model above was specified as

| | $\beta_0$ | $\beta_1$ | $\beta_2$ | $\beta_3$ |
| --- | --- | --- | --- | --- |
| Invalid | 1 | -1 | 0 | 0 |
| Uncued | 1 | 1 | -1 | 0 |
| Valid:Conv5 | 1 | 0 | 1 | 1 |
| Valid:Pixels | 1 | 0 | 0 | -1 |

Therefore, *β*_1_ codes Uncued - Invalid, *β*_2_ codes Valid:Conv5 - Uncued, *β*_3_ codes Valid:Conv5 - Valid:Pixels and *β*_0_ codes the Intercept (average performance). Note that the generalised inverse of this matrix was passed to brms (Venables and Ripley 2002).

Each of these population fixed-effects is offset by the random effects of participant (`subj`) and image (`im_code`). We also assume that the offsets for each fixed effect can be correlated (denoted by the single pipe character |). The model thus estimates:

1. Four fixed-effect coefficients. The coefficients coding Valid:Conv5 – Uncued and Uncued – Invalid constitute the key outcome measures of the study. The final coefficient is the analysis of secondary interest.
2. Eight random-effects standard-deviations (four for each fixed-effect, times two for the two random effects).
3. Twelve correlations (six for each pairwise relationship between the fixed-effects, times two for the two random effects).

**Figure S11.**
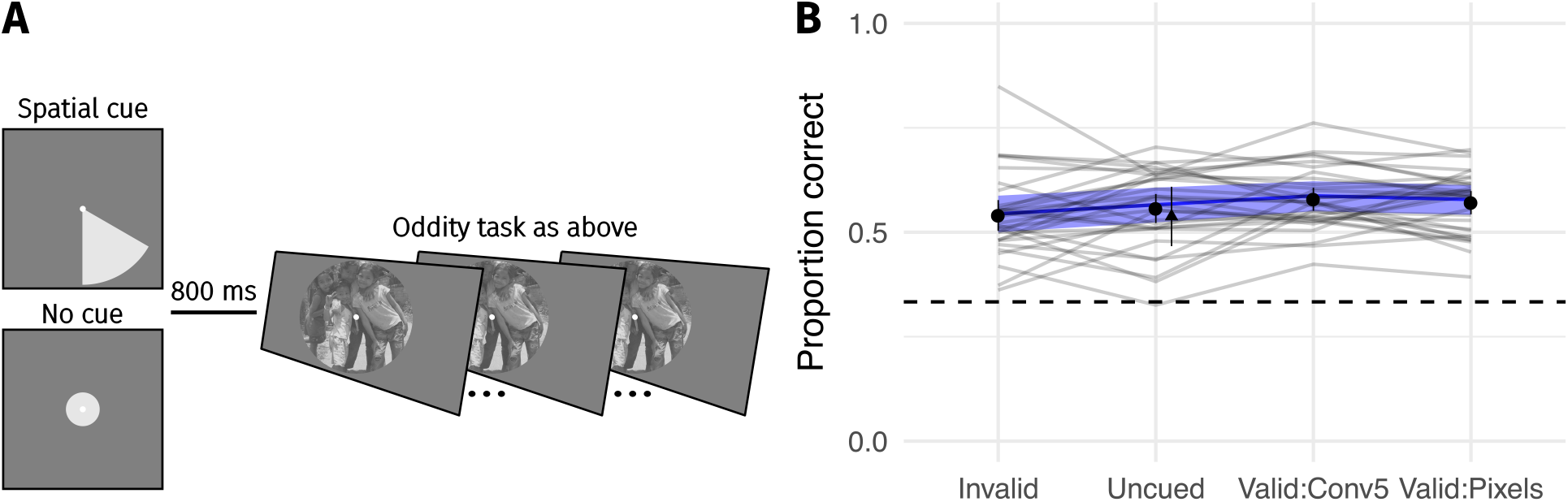
Cueing spatial attention has little effect on performance. **A:** Covert spatial attention was cued to the area of the largest difference between the images (70% of trials; half from conv5 feature MSE; half from pixel MSE) via a wedge stimulus presented before the trial. On 15% of trials the wedge cued an invalid location (smallest pixel MSE), and on 15% of trials no cue was provided (circle stimulus). **B:** Performance as a function of cueing condition for 30 participants. Points show grand mean (error bars show ±2 SE), lines link the mean performance of each observer for each pooling model (based on at least 30 trials; median 65). Blue lines and shaded area show the population mean estimate and 95% credible intervals from the mixed-effects model. Triangle in the Uncued condition replots the average performance from CNN 8 in Figure S8 for comparison.

These parameters were given weakly-informative prior distributions as for Experiment 1 (above): fixed-effects had Cauchy(0, 1) priors, random effect SDs had bounded Cauchy(0.2, 1) priors, and correlation matrices had LKJ(2) priors.

To judge the study outcome we pre-defined a region of practical equivalence (ROPE) around zero effect (0) of [-0.2, 0.2] on the linear predictor scale. This corresponds to odds ratios of [0.82, 1.22]. Our decision rules were then:

- If the 95% credible interval of the parameter value falls outside the ROPE, we consider there to be a credible difference between the conditions.
- If the 95% credible interval of the parameter value falls fully within the ROPE, we consider there to be no practical difference between the conditions. This does not mean that there is no effect, but only that it is unlikely to be large.
- If the 95% credible interval overlaps the ROPE, the data are ambiguous as to the conclusion for our hypothesis. This does not mean that the data give no insight into the direction and magnitude of any effect, but only that they are ambiguous with respect to our decision criteria.

For more discussion of this approach to hypothesis testing, see (Kruschke 2015).

##### 2.4.2 Results and discussion

The results of this experiment are shown in Figure S11B. While mean performance across conditions was in the expected direction for all effects, no large differences were observed. Specifically, the population-level coefficient estimate on the linear predictor scale for the difference between the Valid:Conv5 cueing condition and the uncued condition was 0.09, 95% CI [-0.05, 0.22], p(*β* < 0) = 0.1. Given our decision rules above, the coefficient does not fall wholely within the ROPE and therefore this result is somewhat inconclusive; in general the difference is rather small and so large “true” effects of spatial cueing are quite unlikely. Similarly, we find no large difference between uncued performance and the invalid cues (0.09, 95% CI [-0.07, 0.25], p(*β* < 0) = 0.141). Based on our pre-registered cutoff for a meaningful effect size we conclude that cueing spatial attention in this paradigm results in effectively no performance change.

**Figure S12.**
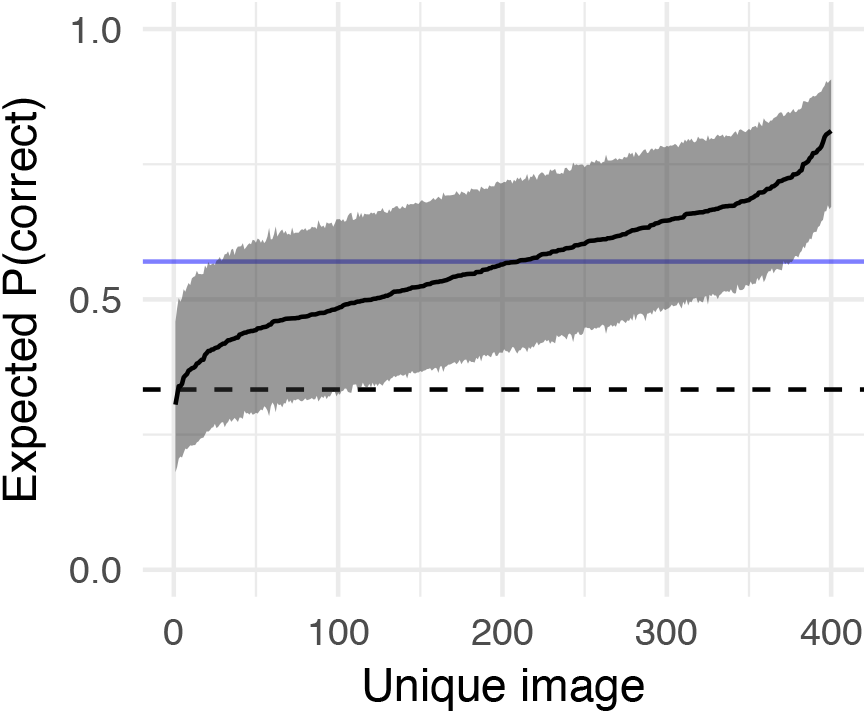
Performance depends strongly on the image within the CNN 8 model (data from Experiment 3). Solid black line links model estimates of each image’s difficulty (the posterior mean of the image-specific model intercept, plotted on the performance scale). Shaded region shows 95% credible intervals. Dashed horizontal line shows chance performance; solid blue horizontal line shows mean performance.

We further hypothesised that the conv5 cue would be more informative (resulting in a larger performance improvement) than the pixel MSE cue. Note that for 269 of 400 images the conv5 and pixel MSE cued the same or neighbouring wedges, meaning that the power of this experiment to detect differences between these conditions is limited. Consistent with this and contrary to our hypothesis, we find no practical difference between the Valid:Conv5 and Valid:Pixels conditions, 0.04, 95% CI [-0.07, 0.14], p(*β* < 0) = 0.253. Note that for this comparison, the 95% credible intervals for the parameter fall entirely within the ROPE, leading us to conclude that there is no practical difference between these conditions in our experiment.

In an additional (exploratory) analysis we assessed whether this experiment also produced evidence for source-image-dependence, consistent with the main experiment (scene-like vs texture-like) and Experiment 2 above. To do so, we plot the image-specific intercepts estimated by the model above. We examine this rather than the raw data because cueing conditions were randomly assigned to each image for each subject, meaning that the mean performance of the images will depend on this randomisation (though, given our results, the effects are likely to be small). The image-specific intercept from the model estimates the difficulty of each image, holding cueing condition constant. While the posterior means for some images were close to chance, and the 95% credible intervals associated with about 100 images overlapped chance performance, approximately 30 images were easily discriminable from their model syntheses, lying above the mean performance for all images with the CNN 8 model (Figure S12). These results again corroborate the evidence above, that the fidelity of appearance matching by the CNN scene appearance model depends substantially on the source image.

To conclude, our results here suggest that if cueing spatial attention improves the “resolution” of the periphery, then the effect is very small. Cohen and colleages (2016) have suggested that an ensemble representation serves to create phenomenal experience of a rich visual world, and that spatial attention can be used to gain more information about the environment beyond simple summary statistics. The results here are contrary to this idea, at least for the specific task and setting we measure here.

Note however that other experimental paradigms may in general be more suitable for assessing the influence of spatial attention than a temporal oddity paradigm. For example, in temporal oddity participants may choose to reallocate spatial attention after the first interval is presented (e.g. on invalid trials pointing at regions of sky). In this respect a single-interval yes-no design (indicating original / synthesis) might be preferable. However, analysis of such data with standard signal detection theory would need to assume that the participants’ decision criteria remain constant over trials, whereas it seems likely that decision criteria would depend strongly on the image. To remain consistent with our earlier experiments we nevertheless employed a three-alternative temporal oddity task here; future work could assess whether our finding of minimal influence of spatial cueing depends on this choice.

#### 2.5 Experiment 4: Sensitivity to local texture distortions

In Figure 3 of the main paper, we presented a demonstration that humans can be quite insensitive to even large texture-like distortions that fall onto texture-like regions of the image. Here we quantify this claim more rigorously by measuring human sensitivity to local texture distortions applied to either scene-like / inhomogenous or texture-like / homogenous regions of images.

##### 2.5.1 Methods

###### 2.5.1.1 Participants

Four observers participated in the experiment; two authors plus two naïve observers paid 10 Euro per hour.

###### 2.5.1.2 Stimuli

As above, we used images from the MIT 1003 database (Judd, Durand, and Torralba 2012; Judd et al. 2009). Authors were familiar with these images; naïve observers were not. We handpicked images that contained local regions that were “scene-like” or “texture-like” (as described in the main manuscript). The regions were choosen in a way that their midpoint has a distance of 128 px (approximately 6 degrees) to the center of the image. In total this resulted in 389 unique images, of which 229 contained a “scene-like” region and 160 images contained a “texture-like” region.

For each image we perturbed a circular patch in the center of the texture/object region. This patch was texturised by using the texture model of Gatys et al (2015). Note that this is the texture-model *not* the CNN-model (using radial and angular pooling regions) reported in the experiments above. For each original image, we generated new images containing distortions of different sizes (radii of 40, 70, 85 and 100 px, corresponding to approximately 2, 3.4, 4.1 and 4.9 degrees). Some example images are shown in Figure S13. In total we therefore used 389 unique images and 389^*^4 synthesised images as stimuli in our experiment.

###### 2.5.1.3 Procedure

Observers discriminated which image contained the distortion in a 2IFC paradigm. Each image was presented for 200~ms with a 1000~ms inter-stimulus interval, after which the observer had 1200~ms to respond. The original, unmodified image could appear either first or second; the other image was the same but contained the circular distortion. Observers fixated a spot (Thaler et al. 2013) in the centre of the screen. Feedback was provided, and eyetracking was not used.

All observers performed 389 trials. To avoid effects of familiarity with the distortion region, each observer saw each original image only once (that is, each original image was randomly assigned to one of the four distortion scales for each observer).

###### 2.5.1.4 Data analysis

We discarded trials for which participants made no response (N = 8), leaving a total of 1548 trials for further analysis.

**Figure S13.**
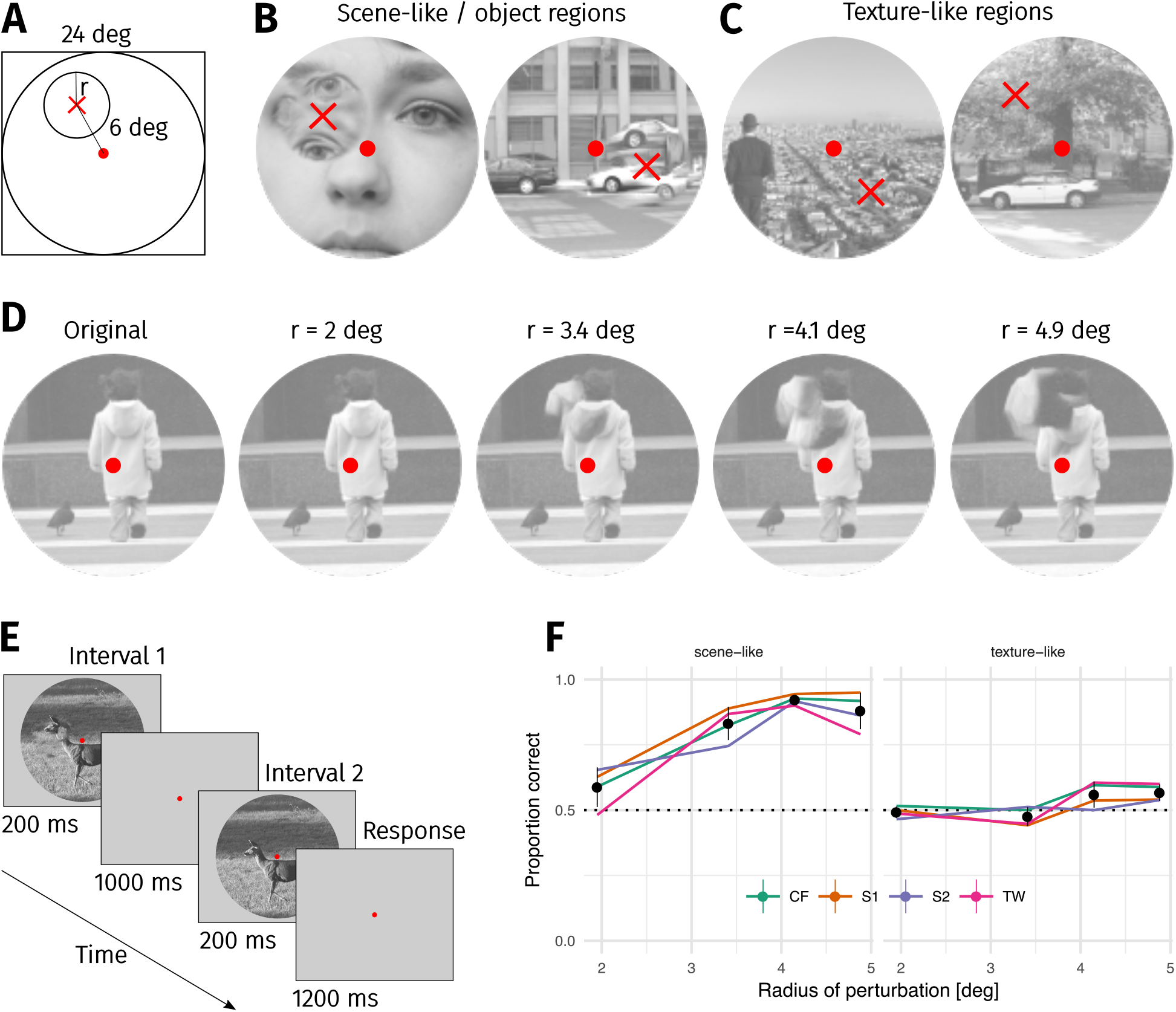
Sensitivity to local texture distortions depends on image content. **A:** A circular patch of an image was replaced with a texture-like distortion. In different experimental conditions the radius of the patch was varied. **B:** Two example images in which a ‘scene-like’ or inhomogenous region is distorted. **C:** Two example images in which a ‘texture-like’ or homogenous region is distorted. **D:** Examples of an original image and the four distortion sizes used in the experiment. **E:** Depiction of the 2IFC task, in which the observer reported whether the first or second image contained the distortion. **F:** Proportion correct as a function of distortion radius in scene-like (left) and texture-like (right) image regions. Lines link the performance of each observer (each point based on a median of 51.5 trials; min 31, max 62). Points show mean of observer means, error bars show ±2 SEM.

##### 2.5.2 Results and discussion

The results of the experiment show that observers can be quite insensitive to even large texture-like distortions that are applied to texture-like image regions. Performance for distortions of nearly 5 degrees radius (i.e. nearly entering the foveal fixation point) was still quiet close to chance. Conversely, distorting scene-like or inhomogenous regions (e.g. containing object borders) is readily detectable for the three largest distortion patch sizes we tested. As shown in Figure 3 of the main paper, the visibility of texture-like distortions mainly depends on the image content to which they are applied.

#### 2.6 Predicting the difficulty of individual images

As shown above, some images are easier than others. We assessed whether an image-based metric considering the difference between original and synthesised images could predict difficulty at the image level. Specifically, we asked whether the mean squared-error (MSE) between the original and synthesised images in two feature spaces (conv5 and pixels) could predict the relative difficulty of the source images. Note that we performed this analysis first on the results of Experiment 1 (Figure S8), and that these results were used to inform the hypothesis regarding the usefulness of conv5 vs pixel cues presented in Experiment 3, above. We subsequently performed the same analysis on the data from Experiment 3. We present both analyses concurrently here for ease of reading, but the reader should be aware of the chronological order.

##### 2.6.1 Methods

We computed the mean squared error between the original and synthesised images in two feature spaces. First, the MSE in the pixel space was used to represent the physical difference at all spatial scales. Second, the difference in feature activations in the conv5 layer of the VGG network was used as an abstracted feature space which may correspond to aspects of human perception (e.g. Kubilius, Bracci, and Op de Beeck 2016). Both are also correlated with the final value of the loss function from our synthesis procedure. As a baseline we fit a mixed-effects logistic regression containing fixed-effects for the levels of the CNN model and a random effect of observer on all fixed effect terms. As a “saturated” model (a weak upper bound) we added a random effect for image to the baseline model (that is, each image is uniquely predicted given the available data). Using the scale defined by the baseline and saturated models, we then compared models in which the image-level predictor (pixel or conv5 MSE, standardised to have zero mean and unit variance within each CNN model level) was added as an additional linear covariate to the baseline model. That is, each image was associated with a scalar value of pixel / conv5 MSE with each synthesis. Additional image-level predictors were compared but are not reported here because they performed similarly or worse than the conv5 or pixel MSE.

As above, we compared the models using the LOOIC information criterion that estimates out-of-sample prediction error on the deviance scale. Qualitatively similar results were found using ten-fold crossvalidation for models fit with penalised maximum-likelihood in lme4.

##### 2.6.2 Results

For the dataset from Experiment 1, the LOOIC favoured the model containing conv5 MSE over the pixel MSE (LOOIC difference 18.2, SE = 8.3) and the pixel MSE over the baseline model (LOOIC difference 25.3, SE = 10.9)—see Figure S14A. The regression weight of the standardised pixel MSE feature fit to all the data was 0.04 (95% credible interval = 0.15–0.07), and the weight of the standardised conv5 feature was 0.04 (0.2–0.11; presented as odds ratios in Figure S14C). Therefore, a one standard deviation increase in the conv5 feature produced a slightly larger increase in the linear predictor (and thus the expected probability) than the pixel MSE, in agreement with the model comparison.

Applying this analysis to the data from Experiment 3 lead to similar results (Figure S14B, d). The LOOIC favoured the model containing conv5 MSE over the pixel MSE (LOOIC difference 49.9, SE = 13.3) and the pixel MSE over the baseline model (LOOIC difference 62.4, SE = 16.2). Note that the worse performance of the image metric models relative to the saturated model (compared to Figure S14A) is because the larger data mass in this experiment provides a better constraint for the random effects estimates of image. The regression weight of the standardised pixel MSE feature fit to all the data was 0.03 (95% credible interval = 0.14–0.08), and the weight of the standardised conv5 feature was 0.03 (0.21–0.15).

**Figure S14.**
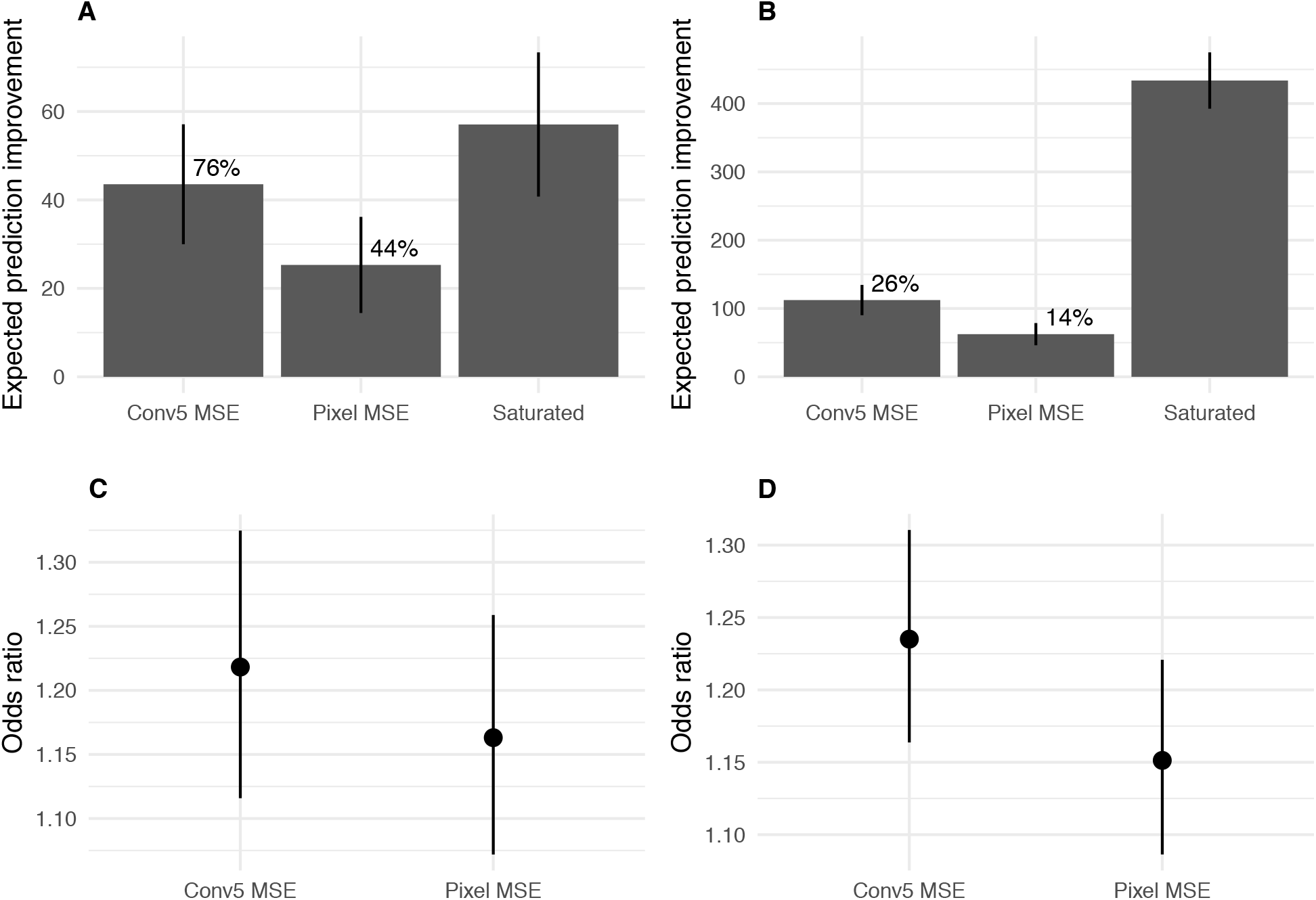
Predicting image difficulty using image-based metrics. **A**: Expected prediction improvement over a baseline model for models fit to the data from Experiment 1 (Figure S8), as estimated by the LOOIC (Vehtari et al., 2016). Values in deviance units (-2 * log likelihood; higher is better). Error bars show ±2 1 SE. Percentages are expected prediction improvement relative to the saturated model. **B**: Same as **A** but for the data from Experiment 3 (Figure S11). **C**: Odds of a success for a one SD increase in the image predictor for data from Experiment 1. Points show mean and 95% credible intervals on odds ratio (exponentiated logistic regression weight). **D**: As for **C** for Experiment 3.

These results show that the difficulty of a given image can be to some extent predicted from the pixel differences or conv5 differences, suggesting these might prove useful full-reference metrics, at least with respect to the distortions produced by our CNN model.

## Footnotes

^1^Many other phenomena also demonstrate striking “failures” of peripheral vision, for example change blindness (Rensink, O’Regan, and Clark 1997; O’Regan, Rensink, and Clark 1999) and inattentional blindness (Mack and Rock 1998), though there is some discussion as to what extent these are distinct from crowding (Rosenholtz 2016).

^2^This selection of images is debateable. In particular some “texture-like” images contain scene-like content. Interestingly, “scene-like” images tend to contain human-made content whereas “texture-like” images tend to contain more natural content, though this was not consciously part of our selection criteria (thanks to Corey Ziemba for pointing this out). We selected these images from a scene database to remain consistent with our other experiments (see Supplementary Material). The fact that we do find differences in critical scaling supports our general argument. See Figure S3 and Supplement for further discussion.

## References

Agaoglu Mehmet N., and Susana T. L. Chung. 2016. “Can (Should) Theories of Crowding Be Unified?” Journal of Vision 16 (15):10. https://doi.org/10.1167/16.15.10.

Ariely Dan. 2001. “Seeing Sets: Representation by Statistical Properties.” Psychological Science 12 (2):157–62.

Arnold, Jeffrey B. 2016. Ggthemes: Extra Themes, Scales and Geoms for ‘Ggplot2’.

Auguie Baptiste. 2016. gridExtra: Miscellaneous Functions for ”Grid” Graphics.

Balas, Benjamin, L Nakano, and Ruth Rosenholtz. 2009. “A Summary-Statistic Representation in Peripheral Vision Explains Visual Crowding.” Journal of Vision 9 (12):13.

Block N. 2013. “Seeing and Windows of Integration.” Thought: A Journal of Philosophy.

Bouma H. 1970. “Interaction Effects in Parafoveal Letter Recognition.” Nature 226 (5241):177–78.

Brainard, David H. 1997. “The Psychophysics Toolbox.” Spatial Vision 10 (4):433–36.

Bürkner, Paul-Christian. 2017. “Brms: An R Package for Bayesian Multilevel Models Using Stan.” Journal of Statistical Software 80 (1):1–28. https://doi.org/10.18637/jss.v080.i01.

Chang, Honghua, and Ruth Rosenholtz. 2016. “Search Performance Is Better Predicted by Tileability Than Presence of a Unique Basic Feature.” Journal of Vision 16 (10):13. https://doi.org/10.1167/16.10.13.

Cohen Michael A., Daniel C. Dennett, and Nancy Kanwisher. 2016. “What Is the Bandwidth of Perceptual Experience?” Trends in Cognitive Sciences 20 (5):324–35. https://doi.org/10.1016/j.tics.2016.03.006.

Cornelissen Frans W., Enno M. Peters, and John Palmer. 2002. “The Eyelink Toolbox: Eye Tracking with MATLAB and the Psychophysics Toolbox.” Behavior Research Methods, Instruments, & Computers 34 (4):613–17. https://doi.org/10.3758/BF03195489.

Craven B. J. 1992. “A Table Of *d*′ forM-Alternative Odd-Man-Out Forced-Choice Procedures.” Perception & Psychophysics 51 (4):379–85.

Croner L. J., and E. Kaplan. 1995. “Receptive Fields of P and M Ganglion Cells Across the Primate Retina.” Vision Research 35 (1):7–24.

Dacey D. M., and M. R. Peterse. 1992. “Dendritic Field Size and Morphology of Midget and Parasol Ganglion Cells of the Human Retina.” Proceedings of the National Academy of Sciences 89 (20):9666–70. https://doi.org/10.1073/pnas.89.20.9666.

Dakin, Steven C, and R.J. Watt. 1997. “The Computation of Orientation Statistics from Visual Texture.” Vision Research 37 (22):3181–92. https://doi.org/10.1016/S0042-6989(97)00133-8.

Deza, Arturo, Aditya Jonnalagadda, and Miguel Eckstein. 2017. “Towards Metamerism via Foveated Style Transfer.” arXiv Preprint arXiv:1705.10041.

Ehinger Krista A., and Ruth Rosenholtz. 2016. “A General Account of Peripheral Encoding Also Predicts Scene Perception Performance.” Journal of Vision 16 (2):13. https://doi.org/10.1167/16.2.13.

Faivre, Nathan, Vincent Berthet, and Sid Kouider. 2012. “Nonconscious Influences from Emotional Faces: A Comparison of Visual Crowding, Masking, and Continuous Flash Suppression.” Frontiers in Psychology 3. https://doi.org/10.3389/fpsyg.2012.00129.

Fischer, Jason, and David Whitney. 2011. “Object-Level Visual Information Gets Through the Bottleneck of Crowding.” Journal of Neurophysiology 106:1389–98.

Francis, Gregory, Mauro Manassi, and Michael H. Herzog. 2017. “Neural Dynamics of Grouping and Segmentation Explain Properties of Visual Crowding.” Psychological Review. https://doi.org/10.1037/rev0000070.

Freeman, Jeremy, and Eero P. Simoncelli. 2011. “Metamers of the Ventral Stream.” Nature Neuroscience 14 (9):1195–1201.

Freeman, Jeremy, and Eero P. Simoncelli. 2013. “The Radial and Tangential Extent of Spatial Metamers.” Journal of Vision 13 (9):573–73. https://doi.org/10.1167/13.9.573.

Freeman, Jeremy, Corey M. Ziemba, David J Heeger, Eero P Simoncelli, and J. Anthony Movshon. 2013. “A Functional and Perceptual Signature of the Second Visual Area in Primates.” Nature Neuro-science 16 (7):974–81.

Gatys L. A., A. S. Ecker, and M. Bethge. 2015. “Texture Synthesis Using Convolutional Neural Networks.” In Advances in Neural Information Processing Systems 28.

Haun Andrew M., Giulio Tononi, Christof Koch, and Naotsugu Tsuchiya. 2017. “Are We Underestimating the Richness of Visual Experience?” Neuroscience of Consciousness 3 (1). https://doi.org/10.1093/nc/niw023.

Herzog Michael H., Bilge Sayim, Vitaly Chicherov, and Mauro Manassi. 2015. “Crowding, Grouping, and Object Recognition: A Matter of Appearance.” Journal of Vision 15 (6).

Hoffman Matthew D., and Andrew Gelman. 2014. “The No-U-Turn Sampler: Adaptively Setting Path Lengths in Hamiltonian Monte Carlo.” Journal of Machine Learning Research 15 (Apr):1593–1623.

Jones, Eric, Travis Oliphant, and Pearu Peterson. 2001. SciPy: Open Source Scientific Tools for Python.

Judd, Tilke, Frédo Durand, and Antonio Torralba. 2012. “A Benchmark of Computational Models of Saliency to Predict Human Fixations.” CSAIL Technical Reports, January.

Judd, Tilke, Krista A. Ehinger, F. Durand, and A. Torralba. 2009. “Learning to Predict Where Humans Look.” In IEEE 12th International Conference on Computer Vision, 2106–13. Kyoto. https://doi.org/10.1109/ICCV.2009.5459462.

Keshvari, Shaiyan, and Ruth Rosenholtz. 2016. “Pooling of Continuous Features Provides a Unifying Account of Crowding.” Journal of Vision 16 (3):39. https://doi.org/10.1167/16.3.39.

Kleiner, M, David H Brainard, and Denis G Pell. 2007. “What’s New in Psychtoolbox-3?” Perception 36 (ECVP Abstract Supplement).

Koenderink, Jan, Matteo Valsecchi, Andrea van Doorn, Johan Wagemans, and Karl Gegenfurtner. 2017. “Eidolons: Novel Stimuli for Vision Research.” Journal of Vision 17 (2):7. https://doi.org/10.1167/17.2.7.

Koffka Kurt. 1935. Principles of Gestalt Psychology. Oxford, UK: Harcourt Brace.

Landy, Michael S. 2013. “Texture Analysis and Perception.” The New Visual Neurosciences (Ed. Werner JS, Chalupa LM), 639–52.

Lettvin, Jerome Y. 1976. “On Seeing Sidelong.” The Sciences 16 (4):10–20. https://doi.org/10.1002/j.2326-1951.1976.tb01231.x.

Long, Bria, Talia Konkle, Michael A. Cohen, and George A. Alvarez. 2016. “Mid-Level Perceptual Features Distinguish Objects of Different Real-World Sizes.” Journal of Experimental Psychology: General 145 (1):95–109. https://doi.org/10.1037/xge0000130.

Mack, Arien, and Irvin Rock. 1998. Inattentional Blindness. Vol. 33. MIT press Cambridge, MA.

Manassi, M, B Sayim, and Michael H. Herzog. 2013. “When Crowding of Crowding Leads to Uncrowding.” Journal of Vision 13 (13):10.

Movshon J. Anthony, and Eero P. Simoncelli. 2014. “Representation of Naturalistic Image Structure in the Primate Visual Cortex.” Cold Spring Harbor Symposia on Quantitative Biology 79:115–22. https://doi.org/10.1101/sqb.2014.79.024844.

Neri Peter. 2017. “Object Segmentation Controls Image Reconstruction from Natural Scenes.” Edited by Christopher C. Pack. PLOS Biology 15 (8):e1002611. https://doi.org/10.1371/journal.pbio.1002611.

Okazawa, Gouki, Satohiro Tajima, and Hidehiko Komatsu. 2015. “Image Statistics Underlying Natural Texture Selectivity of Neurons in Macaque V4.” Proceedings of the National Academy of Sciences 112 (4):E351–E360. https://doi.org/10.1073/pnas.1415146112.

O’Regan, J Kevin, Ronald A Rensink, and James J Clar. 1999. “Change-Blindness as a Result of ‘Mudsplashes’.” Nature 398 (6722):34–34.

Parkes, L, J Lund, A Angelucci, JA Solomon, and M Morga. 2001. “Compulsory Averaging of Crowded Orientation Signals in Human Vision.” Nature Neuroscience 4 (7):739–44. https://doi.org/10.1038/89532.

Pelli, Denis G. 1997. “The VideoToolbox Software for Visual Psychophysics: Transforming Numbers into Movies.” Spatial Vision 10 (4):437–42.

Pelli, Denis G, and K A Tillma. 2008. “The Uncrowded Window of Object Recognition.” Nature Neuroscience 11 (10):1129–35.

Portilla, J, and Eero P Simoncell. 2000. “A Parametric Texture Model Based on Joint Statistics of Complex Wavelet Coefficients.” International Journal of Computer Vision 40 (1):49–70.

R Core Team. 2017. R: A Language and Environment for Statistical Computing. Vienna, Austria: R Foundation for Statistical Computing.

Rensink, Ronald A, J Kevin O’Regan, and James J. Clark. 1997. “To See or Not to See: The Need for Attention to Perceive Changes in Scenes.” Psychological Science 8 (5):368–73.

Rosen S., R. Chakravarthi, and Denis G Pell. 2014. “The Bouma Law of Crowding, Revised: Critical Spacing Is Equal Across Parts, Not Objects.” Journal of Vision 14 (6):10–10. https://doi.org/10.1167/14.6.10.

Rosenholtz Ruth. 2016. “Capabilities and Limitations of Peripheral Vision.” Annual Review of Vision Science 2 (1):437–57. https://doi.org/10.1146/annurev-vision-082114-035733.

Rosenholtz, Ruth, Jie Huang, and Krista A. Ehinger. 2012. “Rethinking the Role of Top-down Attention in Vision: Effects Attributable to a Lossy Representation in Peripheral Vision.” Frontiers in Psychology 3:13.

Rosenholtz, Ruth, Jie Huang, Alvin Raj, Benjamin Balas, and Livia Ilie. 2012. “A Summary Statistic Representation in Peripheral Vision Explains Visual Search.” Journal of Vision 12 (4).

Saarela, T P, B Sayim, Gerald Westheimer, and Michael H. Herzog. 2009. “Global Stimulus Configuration Modulates Crowding.” Journal of Vision 9 (2):5.

Seth, Anil K. 2014. “A Predictive Processing Theory of Sensorimotor Contingencies: Explaining the Puzzle of Perceptual Presence and Its Absence in Synesthesia.” Cognitive Neuroscience 5 (2):97–118. https://doi.org/10.1080/17588928.2013.877880.

Stan Development Team. 2017. “Stan: A C++ Library for Probability and Sampling, Version 2.14.0.”.

Thaler, L, A C Schütz, M A Goodale, and K R Gegenfurtne. 2013. “What Is the Best Fixation Target? The Effect of Target Shape on Stability of Fixational Eye Movements.” Vision Research 76:31–42.

Vickery T. J., W. M. Shim, R. Chakravarthi, Y. V. Jiang, and R. Luedema. 2009. “Supercrowding: Weakly Masking a Target Expands the Range of Crowding.” Journal of Vision 9 (February):12–12. https://doi.org/10.1167/9.2.12.

Wallis, Thomas S. A., and Peter J Be. 2012. “Image Correlates of Crowding in Natural Scenes.” Journal of Vision 12 (7):1–19. https://doi.org/10.1167/12.7.6.

Wallis, Thomas S. A., Matthias Bethge, and Felix A. Wichmann. 2016. “Testing Models of Peripheral Encoding Using Metamerism in an Oddity Paradigm.” Journal of Vision 16 (2):4. https://doi.org/10.1167/16.2.4.

Wallis, Thomas S. A., Christina M. Funke, Alexander S. Ecker, L. A. Gatys, Felix A. Wichmann, and Matthias Bethge. 2017. “A Parametric Texture Model Based on Deep Convolutional Features Closely Matches Texture Appearance for Humans.” Journal of Vision 17 (12):5. https://doi.org/10.1167/17.12.5.

Wandell, Brian A. 1995. Foundations of Vision. Sinauer Associates.

Watson, Andrew B. 2014. “A Formula for Human Retinal Ganglion Cell Receptive Field Density as a Function of Visual Field Location.” Journal of Vision 14 (7):15:1–17. https://doi.org/10.1167/14.7.15.

Whitney, David, J Haberman, and TD Sweeny. 2014. “From Textures to Crowds: Multiple Levels of Summary Statistical Perception.” The New Visual Neurosciences, 695–710.

Wickham Hadley. 2009. Ggplot2: Elegant Graphics for Data Analysis. New York: Springer.

Wickham Hadley.. 2011. “The Split-Apply-Combine Strategy for Data Analysis.” Journal of Statistical Software 40 (1):1–29.

Wickham, Hadley, and Romain Francois. 2016. Dplyr: A Grammar of Data Manipulation.

Xie Yihui. 2013. “Knitr: A Comprehensive Tool for Reproducible Research in R. BT - Implementing Reproducible Computational Research.” In Implementing Reproducible Computational Research, edited by Victoria Stodden, Frederich Leisch, and Roger D Pen. Chapman & Hall/CRC.

Xie Yihui.. 2015. Dynamic Documents with R and Knitr, Second Edition. 2nd ed. Chapman & Hall/CRC.

Zhang, Xuetao, Jie Huang, Serap Yigit-Elliott, and Ruth Rosenholtz. 2015. “Cube Search, Revisited.” Journal of Vision 15 (3):9. https://doi.org/10.1167/15.3.9.

Ziemba Corey M., Jeremy Freeman, J. Anthony Movshon, and Eero P. Simoncelli. 2016. “Selectivity and Tolerance for Visual Texture in Macaque V2.” Proceedings of the National Academy of Sciences 113 (22):E3140–E3149. https://doi.org/10.1073/pnas.1510847113.

## References

Freeman, Jeremy, and Eero P. Simoncelli. 2011. “Metamers of the Ventral Stream.” Nature Neuroscience 14 (9):1195–1201.

Gelman, Andrew, Jessica Hwang, and Aki Vehtari. 2014. “Understanding Predictive Information Criteria for Bayesian Models.” Statistics and Computing 24 (6):997–1016. https://doi.org/10.1007/s11222-013-9416-2.

Kruschke John. 2015. Doing Bayesian Data Analysis: A Tutorial with R, JAGS, and Stan. Academic Press.

Kubilius, Jonas, Stefania Bracci, and Hans P. Op de Beeck. 2016. “Deep Neural Networks as a Computational Model for Human Shape Sensitivity.” PLoS Comput Biol 12 (4):e1004896.

Macmillan, N A, and C D Creelman. 2005. Detection Theory: A User’s Guide. Mahwah, NJ: Lawrence Erlbaum.

McElreath Richard. 2016. Statistical Rethinking: A Bayesian Course with Examples in R and Stan. Texts in Statistical Science 122. Boca Raton London New York: CRC Press, Taylor & Francis Group.

Simonyan, Karen, and Andrew Zisserman. 2015. “Very Deep Convolutional Networks for Large-Scale Image Recognition.” ICLR abs/1409.1556.

Stan Development Team. 2015. Stan Modeling Language Users Guide and Reference Manual, Version 2.10.0.

Stan Development Team. 2017. “Stan: A C++ Library for Probability and Sampling, Version 2.14.0.”.

Vehtari, Aki, Andrew Gelman, and Jonah Gabry. 2016. “Practical Bayesian Model Evaluation Using Leave-One-Out Cross-Validation and WAIC_ú_.” arXiv Preprint arXiv:1507.04544.

Venables W. N., and B. D. Ripley. 2002. Modern Applied Statistics with S. Fourth. New York: Springer.

